# An Acentriolar Centrosome At The *C. elegans* Ciliary Base

**DOI:** 10.1101/2020.08.21.260547

**Authors:** Joachim Garbrecht, Triin Laos, Elisabeth Holzer, Margarita Dillinger, Alexander Dammermann

**Author notes:** Contributed equally (Joachim Garbrecht, Triin Laos). Lead contact: Alexander Dammermann, Max Perutz Labs, Dr Bohr-Gasse 9/4, A-1030 Vienna, Austria.

## Abstract

In animal cells the functions of the cytoskeleton are coordinated by centriole-based centrosomes via microtubule-nucleating γ-tubulin complexes embedded in the pericentriolar material or PCM [1]. PCM assembly has been best studied in the context of mitosis, where centriolar SPD-2 recruits PLK-1, which in turn phosphorylates key scaffolding components such as SPD-5 and CNN to promote expansion of the PCM polymer [2–4]. To what extent these mechanisms apply to centrosomes in interphase or in differentiated cells remains unclear [5]. Here, we examine a novel type of centrosome found at the ciliary base of *C. elegans* sensory neurons, which we show plays important roles in neuronal morphogenesis, cellular trafficking and ciliogenesis. These centrosomes display similar dynamic behavior to canonical, mitotic centrosomes, with a stable PCM scaffold and dynamically localized client proteins. Unusually, however, they are not organized by centrioles, which degenerate early in terminal differentiation [6]. Yet, PCM not only persists but continues to grow with key scaffolding proteins including SPD-5 expressed under control of the RFX transcription factor DAF-19. This assembly occurs in the absence of the mitotic regulators SPD-2, AIR-1 and PLK-1, but requires tethering by PCMD-1, a protein which also plays a role in the initial, interphase recruitment of PCM in early embryos [7]. These results argue for distinct mechanisms for mitotic and non-mitotic PCM assembly, with only the former requiring PLK-1 phosphorylation to drive rapid expansion of the scaffold polymer.

**ETOC BLURB:** Centrioles play a critical role in mitotic centrosome assembly. Here, Garbrecht *et al.* show that pericentriolar material (PCM) persists at the ciliary base of *C. elegans* sensory neurons after centriole degeneration, where it contributes to neuronal morphogenesis and cellular trafficking. Remarkably, this PCM displays dynamic properties similar to canonical centrosomes, yet its continued assembly and maintenance is independent of known mitotic regulators, suggesting differential mechanisms for mitotic and non-mitotic centrosome assembly.

**HIGHLIGHTS:**

- PCM persists at the acentriolar ciliary base in *C. elegans*
- PCM assembles in a SPD-2, AIR-1 and PLK-1-independent manner
- PCMD-1 tethers PCM at the ciliary base in the absence of centrioles
- PCM is required for neuronal morphogenesis and cilium assembly

## RESULTS AND DISCUSSION

The nematode *C. elegans* represents perhaps one of the best understood experimental models for mitotic centrosome assembly. A key feature of *C. elegans* is that the architecture of the syncytial gonad makes it possible to use RNAi to generate oocytes whose cytoplasm is reproducibly depleted of targeted gene products via a process that does not depend on intrinsic protein turnover. Fertilization then triggers embryos to attempt their first cell division in the absence of the target protein, yielding clear and distinctive phenotypes, which have aided in the identification of the core machinery for centriole assembly, as well as proteins required for PCM assembly and function [8]. The resultant comprehensive parts list has also inspired efforts to reconstitute these processes *in vitro* [4, 9–11]. Central to mitotic PCM assembly is the coiled coil protein SPD-5, which forms a polymeric scaffold around centrioles (Figure 1A) [4, 12]. PCM ‘client’ proteins including the nucleator γ-tubulin, microtubule regulators such as TAC-1 and ZYG-9 as well as tubulin itself either dynamically concentrate within the PCM or stably bind to the polymer lattice [11, 13]. SPD-5 spontaneously self-assembles *in vitro*, a process potentiated by SPD-2 and the polo-like kinase PLK-1 [4]. *In vivo*, this process is directed to occur around centrioles with centriole-localized SPD-2 recruiting both PLK-1 and its activator Aurora A/AIR-1 [2]. Laser ablation experiments have shown that centrioles are essential for mitotic PCM growth, although existing PCM polymer can be maintained by PLK-1 acting within the PCM to counter the destabilizing effect of phosphatases such as PP2A [2, 14]. Centrioles are further essential for PCM structural integrity, potentially by acting as anchoring sites for tethering proteins such as PCMD-1, a putative ortholog of pericentrin [2, 7].

**Figure 1:**
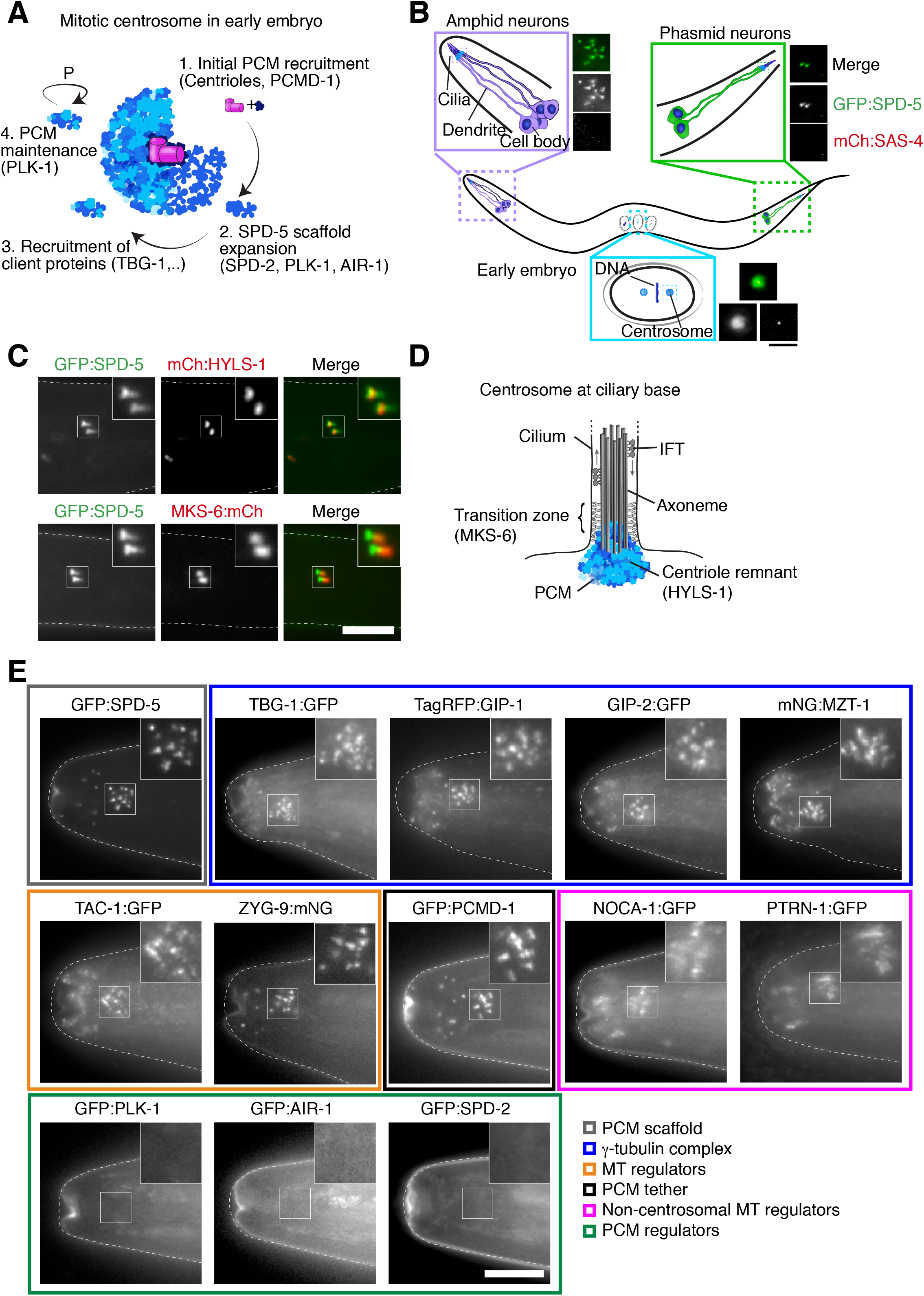
PCM is maintained at the *C. elegans* ciliary base in the absence of centrioles and mitotic regulators. (**A**) Schematic illustrating steps in assembly and maintenance of the mitotic centrosome: (1) Initial, interphase recruitment of PCM to centrioles in a manner dependent on the putative pericentrin ortholog PCMD-1. (2) SPD-2, AuroraA/AIR-1 and PLK-1-dependent expansion of the SPD-5 PCM scaffold. (3) Recruitment of PCM client proteins including γ-tubulin, TAC-1, etc.. (4) Maintenance of the PCM scaffold via PLK-1 acting within the PCM. (**B**) Schematic of adult worm illustrating positions of amphid and phasmid sensory neurons in the head and tail of the animal, and developing embryos. Images show PCM (GFP:SPD-5, green) and centriole (mCherry:SAS-4, red) signal at the ciliary base and early embryonic centrosome. SPD-5 signal persists in ciliated neurons despite loss of centrioles. (**C**) Localization of PCM (GFP:SPD-5, green) relative to remnant of degenerated basal body (mCherry:HYLS-1, red) and ciliary transition zone (MKS-6:mCherry, red). PCM signal colocalizes with HYLS-1 but also extends into core of the transition zone. (**D**) Schematic depiction of PCM organization at ciliary base. (**E**) Presence of different functional classes of PCM proteins and non-centrosomal microtubule regulatory proteins (NOCA-1, PTRN-1) at amphid ciliary base revealed using endogenously tagged/endogenous promoter fluorescent fusions as indicated. PCM composition is similar to canonical centrosomes except for the presence of NOCA-1 and PTRN-1 and the lack of the regulators of mitotic PCM assembly, SPD-2, AIR-1 and PLK-1. Scale bars are 5*μ*m. (B, C), 10*μ*m (E). See also Figure S1.

### An acentriolar PCM at the *C. elegans* ciliary base

*C. elegans* is well known for its stereotypical pattern of development, with precisely 959 somatic cells in the adult hermaphrodite [15]. Less widely appreciated is that centrioles and centrosomes are lost in most of those cells, potentially reflecting their lack of further proliferative potential [16]. This is also true for ciliated sensory neurons, where centrioles degenerate soon after terminal cell division, a phenomenon that has helped define their functional contribution to the process of cilium biogenesis [6, 17]. Centriolar structural proteins including SAS-4, SAS-5 and SAS-6 are likewise lost [6]. Yet, surprisingly, γ-tubulin has been shown to persist at the ciliary base in adult worms [18, 19]. This accumulation could simply be a consequence of γ-tubulin capping the free minus ends of ciliary microtubules [20]. However, further examination showed that γ-tubulin is not alone. Endogenously GFP-tagged SPD-5 similarly forms discrete foci in ciliated neurons, including the amphid and phasmid sensory organs in the head and tail of the animal, respectively, which are the subject of most studies on cilia in *C. elegans* (Figure 1B, [21]). SPD-5 foci are also observed in cephalic and labial neurons, visible as a cluster of puncta surrounding the mouth of the worm, as well as in the sensory neurons of the male tail (Figure S1A). These foci, which are similar in size to interphase centrosomes in the early embryo, are all the more prominent given the near absence of SPD-5 signal in other somatic tissues of the adult worm. Closer inspection shows this SPD-5 accumulation to be in the region of the degenerated basal body and extending into the ciliary transition zone, marked by HYLS-1 and MKS-6 [6, 22], respectively (Figure 1C, D). Thus, a type of PCM remains following centriole degeneration in the late-stage embryo and persists throughout the lifetime of the animal.

Besides the scaffolding protein SPD-5 and the nucleator γ-tubulin (TBG-1 in *C. elegans*), the PCM in the early embryo is characterized by the presence of numerous client proteins that regulate microtubule nucleation, stability and/or organization. These include other members of the γ-tubulin complex, the GCP-2 homolog GIP-2, the GCP-3 homolog GIP-1 and Mozart/MZT-1 [23, 24] and the microtubule polymerising complex of TACC/TAC-1 and ch-TOG/ZYG-9 [25]. All of these proteins are also found in ciliated neurons (Figure 1E), consistent with the reported presence of a microtubule-organizing center at the dendritic tip [19]. We could also detect clear foci of the putative homolog of pericentrin, PCMD-1, which helps recruit/stabilize PCM around centrioles in the early embryo [7]. Also present at the ciliary base are the microtubule anchoring protein Ninein/NOCA-1 and the minus end stabilizer CAMSAP/PTRN-1, which in *C. elegans* are not found at the mitotic centrosome in the early embryo but have been reported to localize to non-centrosomal microtubule organizing centers in the larval and adult skin, germline and intestine (Figure S1B) [24, 26]. Another striking difference to mitotic centrosomes is the complete absence of the PCM regulators SPD-2, AIR-1 and PLK-1 [2, 4, 27–29], none of which could be detected at the ciliary base in contrast to their prominent localization to centrioles and/or the PCM in early embryos (Figure S1C). There are then several critical differences between the neuronal PCM and the canonical, mitotic PCM that has been the subject of most *C. elegans* studies so far.

### SPD-5 is specifically expressed in ciliated neurons under control of the RFX transcription factor DAF-19

Unlike other microtubule-organizing centers in the adult worm the acentriolar centrosome in ciliated neurons derives from a canonical, centriole-organized centrosome present at the initiation of ciliogenesis in the late-stage embryo [6]. A potential explanation for the persistence of PCM is that there is no counteracting phosphatase driving PCM polymer disassembly, as there is during mitosis [2, 14]. The PCM at the ciliary base could then simply be a remnant left over from the terminal cell division, surviving the loss of its core organizing structure, the centriole, and the kinase that drove its assembly, PLK-1. However, further analysis showed this is not the case. Thus, our initial attempts to deplete neuronal SPD-5 using the GFP nanobody-directed ZIF-1 degron system [26] expressed under a Pdyf-7 promoter active early during neuronal differentiation were unsuccessful despite the same degron efficiently targeting other proteins [30] (Figure 2A, B). Switching to a heat shock inducible promoter (Phsp-16.41) revealed the reason why: GFP:SPD-5 signal was initially depleted but eventually recovered even after repeated cycles of heat shock-induced degradation (Figure 2C, S2A). SPD-5 therefore continues to be expressed in ciliated neurons. Expression of ciliary genes including components of the transition zone and intraflagellar transport (IFT) machinery in the worm is regulated by the RFX transcription factor DAF-19, which binds to specific X-box elements in the promoter of target genes [31]. A closer inspection of the *spd-5* promoter revealed a predicted X-box near the transcription start site closely matching the consensus sequence [32] (Figure 2D). Consistent with SPD-5 being under the control of DAF-19, *daf-19* mutant animals indeed displayed reduced levels of SPD-5 in sensory neurons (Figure S2B). However, loss of this master regulator of ciliogenesis and neuronal morphogenesis could perturb SPD-5 localization indirectly, via perturbing the organization of the dendritic tip. To test DAF-19 dependency more directly, we mutated the X-box element in the *spd-5* promoter. This did not affect SPD-5 levels in the early embryo (Figure S2C), nor embryonic viability (100%, n=545 embryos vs parental GFP:SPD-5 strain 99.9%, n=709). However, signal in ciliated neurons was strongly reduced (Figure 2E, F). When combined with degron-mediated degradation targeting protein inherited from previous mitotic divisions we were able to efficiently deplete SPD-5 using GFP nanobody-directed ZIF-1 expressed under either tissue-specific or inducible promoters (Figure 2G, H, 3C, D). Thus, SPD-5 is continuously expressed in ciliated neurons and incorporates into an acentriolar PCM in the absence of the regulators of mitotic PCM assembly, SPD-2, AIR-1 and PLK-1.

**Figure 2:**
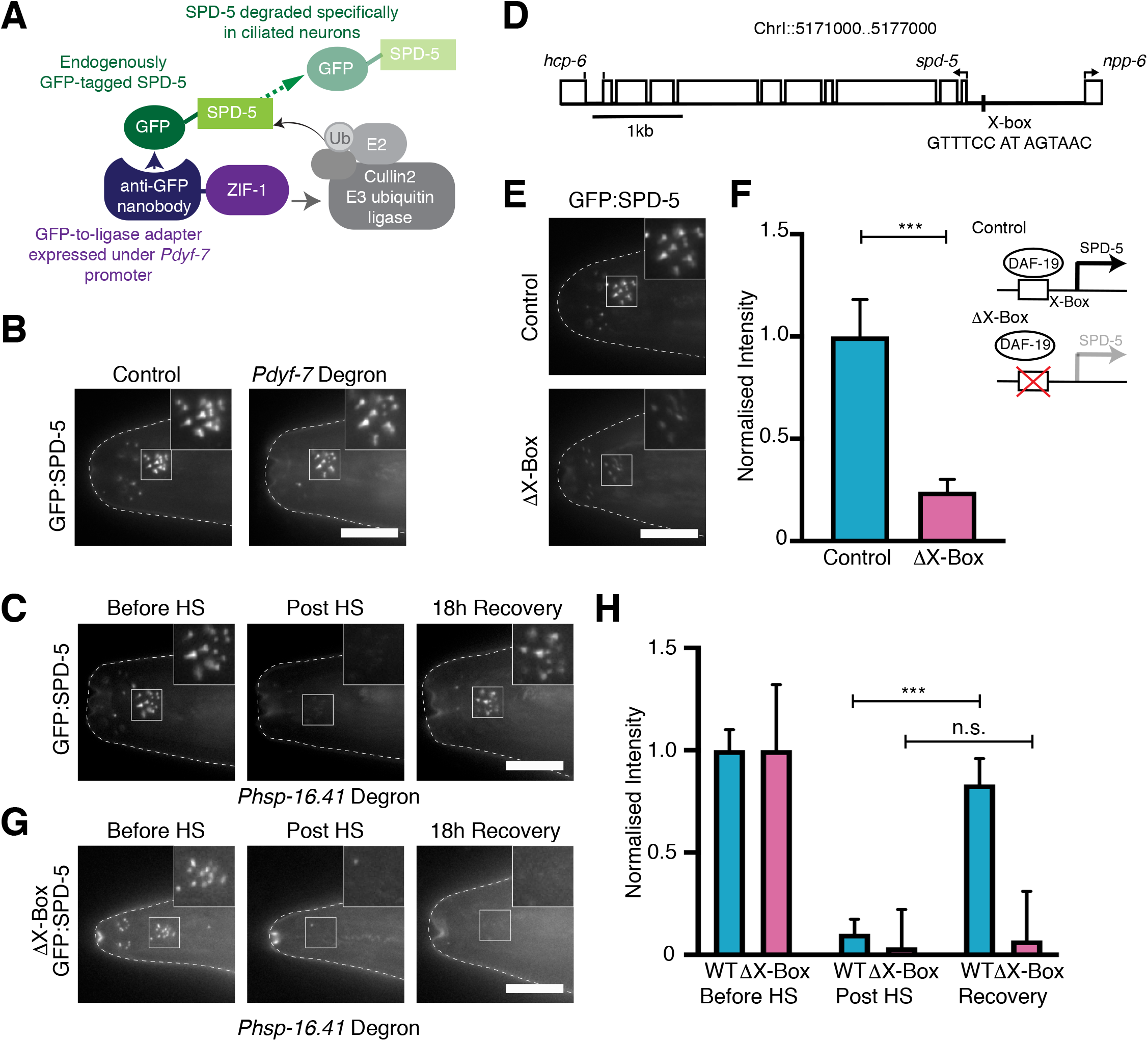
SPD-5 is continuously expressed in ciliated neurons under control of the RFX transcription factor DAF-19. (**A**) Schematic of GFP nanobody-directed ZIF-1 degron system, targeting endogenously GFP-tagged proteins for ubiquitin-mediated degradation. Depending on the promoter used to drive ZIF-1 expression, protein degradation may be tissue-specific (*Pdyf-7* ciliated neurons from comma stage, *Posm-6* from 3-fold stage) or inducible (*Phsp-16.41*, heat shock-inducible). (**B**) *Pdyf-7*-promoter-driven ZIF-mediated degradation fails to result in appreciable decrease in ciliary SPD-5 signal. (**C**) *Phsp-16.41* heat shock promoter-driven ZIF-mediated degradation transiently depletes ciliary SPD-5 signal. However, signal recovers within 18h due to new protein expression. (**D**) Analysis of *spd-5* promoter reveals sequence motif 181bp upstream of ATG closely matching the consensus for DAF-19-regulated X-box genes (GTHNYY AT RRNAAC, [32]). (**E**) Mutation of X-box motif results in a significant decrease in SPD-5 signal at the ciliary base. (**F**) Quantitation of SPD-5 signal based on images as in (E). n=14 animals control, 10 ΔX-box. (**G**) Combining X-box mutation with heat shock promoter-driven ZIF-mediated degradation results in a permanent loss of ciliary SPD-5. (**H**) Quantitation of heat shock degradation experiments shown in (C) and (G). Pre-heat shock signal independently normalized to 1 for wild-type and ΔX-box promoter strains. n=13-17 animals wild-type, 6-15 ΔX-box. Scale bars are 10*μ*m. Error bars are SD. *** t-test, P<0.001, n.s. not significant. See also Figure S2.

### SPD-5 at the ciliary base forms a stable scaffold for the recruitment of client proteins

In the context of the early embryo, SPD-5 forms a polymeric scaffold around centrioles that exhibits little or no exchange of subunits with the surrounding cytoplasm [4, 13]. Certain PCM client proteins, most notably the γ-tubulin complex, stably bind to the SPD-5 lattice and likewise exhibit little cytoplasmic exchange, while others such as TAC-1 and ZYG-9 are more dynamically localized [11, 13]. To examine the dynamic behavior of PCM proteins at the ciliary base we performed fluorescence recovery after photobleaching (FRAP) experiments on live anaesthetized animals, focusing on the phasmids, where a pair of cilia is found in isolation, bleaching one or both PCM foci and comparing the kinetics of recovery, if any, with that at mitotic centrosomes in the early embryo. As can be seen in Figure 3A and B, SPD-5 exhibited essentially no recovery within the timeframe of the experiment. Neither did the γ-tubulin complex protein GIP-2. In contrast, PCM signal of TAC-1 recovered relatively rapidly. While we cannot exclude that recovery kinetics are affected by the geometry of the extended dendrite and the lack of a substantial cytoplasmic pool in differentiated neurons, these results are qualitatively and quantitatively similar to those obtained for canonical centrosomes in the early embryo (Figure S3A, B). This is perhaps not too surprising, given that the dynamic enrichment of certain client proteins within the SPD-5 scaffold has been reproduced with purified proteins *in vitro* [11], indicating that it is an intrinsic property of the scaffold and client proteins themselves. Nevertheless, it argues for a similar type of PCM scaffold being present at the ciliary base and canonical centrosomes.

**Figure 3:**
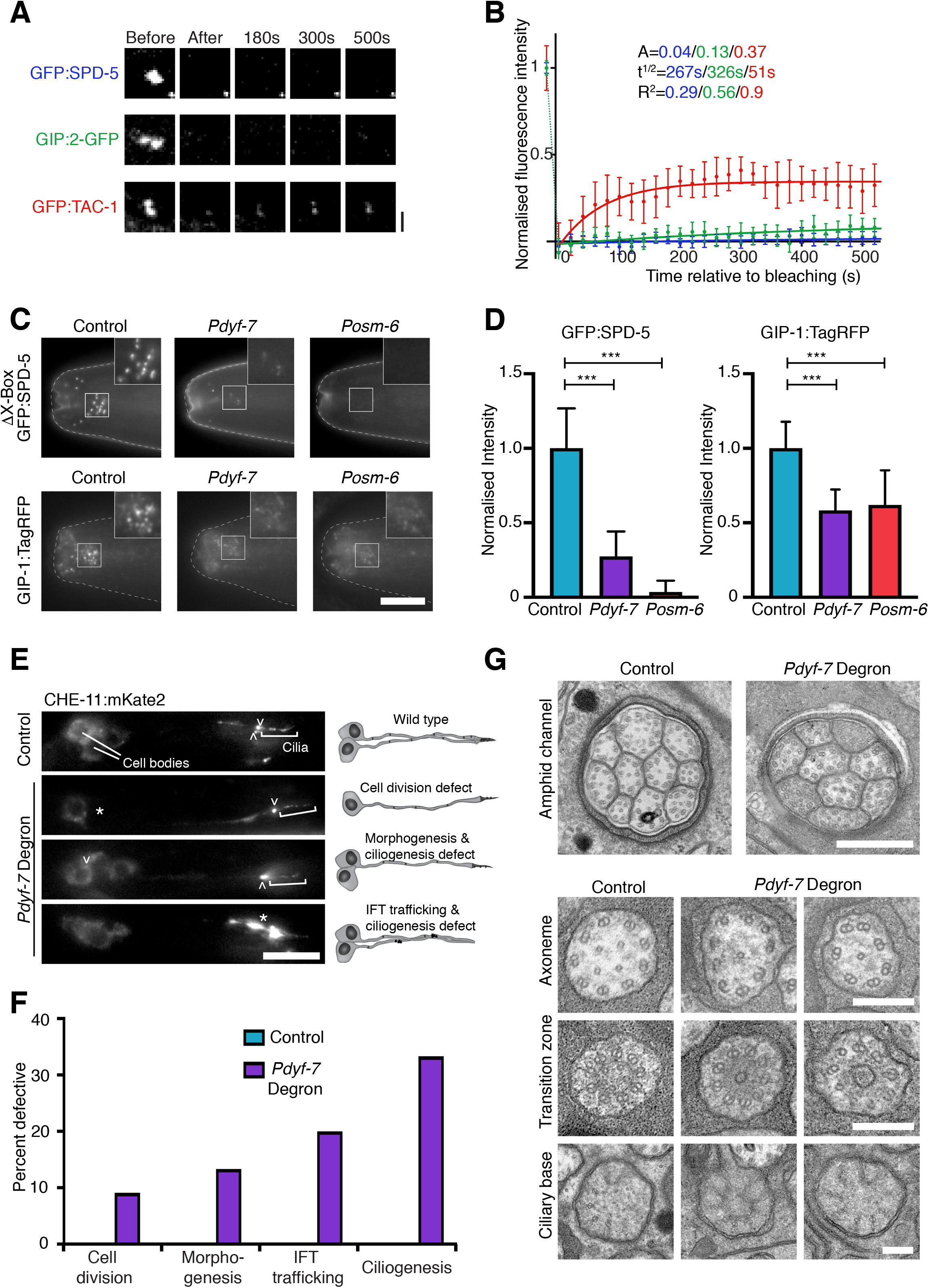
The centrosome at the ciliary base contributes to neuronal morphogenesis and cilium assembly. (**A**, **B**) Fluorescence recovery after photobleaching (FRAP) analysis for PCM proteins at the ciliary base of phasmid neurons. Images (A) and quantitations (B) for FRAP experiments performed on strains expressing GFP:SPD-5 (n=7 animals), GIP-2:GFP (n=8) and GFP:TAC-1 (n=7). As at mitotic centrosomes, SPD-5 forms a PCM scaffold to which the γ-tubulin complex protein GIP-2 is stably bound, while TAC-1 remains in dynamic exchange with the cytoplasmic pool. (**C**, **D**) Tissue-specific degradation of SPD-5 early (*Pdyf-7*) or late in neuronal differentiation (*Posm-6*) results in a loss of GIP-1 from the ciliary base. Images (C) and quantifications (D) for strains expressing ΔX-box GFP:SPD-5 or GIP-1:TagRFP alone (n=21/19 animals) or ΔX-box GFP:SPD-5/ΔX-box GFP:SPD-5 and GIP-1:TagRFP in combination with *Pdyf-7* (n=17/23) or *Posm-6* (n=15/22) promoter-driven GFP-nanobody degrons. (**E**, **F**) Range of defects observed using CHE-11:mKate2 IFT reporter following *Pdyf-7* promoter-driven degradation of SPD-5 in phasmid neurons. Defects include an accumulation of IFT particles along dendrites and abnormal neuronal morphology. Both phenotypes were accompanied by a failure of cilium extension. A number of animals also displayed missing phasmid cell bodies, potentially due to a failure of terminal cell division (n=14 animals control, 33 *Pdyf-7* degron). (**G**) Transmission electron micrographs of amphid cilia of wild-type and *Pdyf-7* degron SPD-5-depleted animals. Cross sectional views reveal missing cilia in the amphid channel (10 cilia extending into the channel in wild-type), missing and incomplete doublet microtubules in the proximal segment of the axoneme and, rarely, disorganized ciliary transition zones. The centriolar remnant previously referred to as transitional fibers located below the transition zone is apparently unaffected. Defects are quantitated in Figure S3G. Scale bars are 2*μ*m (A), 10*μ*m (C, E), 500nm (G, amphid channel), 200nm (G, others). Error bars are 90% CI (B), SD (D). *** t-test, P<0.001. See also Figure S3.

At mitotic centrosomes, SPD-5 is absolutely essential for PCM assembly. Embryos depleted of SPD-5 by RNAi accumulate no detectable PCM around centrioles and spindle assembly completely fails [12]. Testing SPD-5 function in ciliated neurons is complicated by its essential role in prior mitotic cycles, the lack of a robust systemic response to RNAi in *C. elegans* neurons [33] and the low rate of turnover of cytoskeleton proteins. However, the combination of X-box mutation and degron-mediated degradation of protein inherited from the last cell division using a GFP nanobody ZIF-1 fusion expressed either early (Pdyf-7 promoter-driven) or late (Posm-6) during neuronal differentiation efficiently removed SPD-5 from the dendritic tip (Figure 3C, D). SPD-5 degradation resulted in a pronounced loss of the γ-tubulin complex protein GIP-1 from the ciliary base, although the protein remained diffusely localized to the dendritic tip (Figure 3C, D, S3C). Thus, SPD-5 is required to maintain a coherent PCM at the ciliary base, as would be expected for a scaffolding protein.

### The centrosome at the ciliary base contributes to neuronal morphogenesis and cilium assembly

The continued expression of SPD-5 and maintenance of a centrosomal microtubule-organizing center specifically in ciliated neurons suggests that it plays an important role in their development and/or function. A notable feature of amphid and phasmid neurons in the worm is their unique mode of dendrite elongation by a process of retrograde extension, with the cell bodies migrating backwards while the tips remain anchored in place by interactions with the surrounding extracellular matrix [34]. This anchoring has been shown to involve structures at the dendritic tip, in particular the ciliary transition zone [35], but a centrosomal microtubule-organizing center could conceivably also contribute. However, a PCM focus is also observed in the ciliated PQR neuron, which is known to form its dendrite in the canonical, anterograde manner by growth cone extension [36] (Figure S1A). Another potential function concerns the organization of microtubules within neuronal projections. The extended nature of dendrites and axons places particular demands on intracellular transport. To facilitate such transport, microtubules within dendrites and axons generally have a uniform polarity, in the case of axons with their plus ends facing outwards, while microtubules within the shorter dendrites tend to be of mixed polarity or oriented with their minus end facing outwards [37]. In *C. elegans*, dendrites of both ciliated and non-ciliated neurons have microtubules with minus end out polarity. However, ciliated neurons display a higher density of microtubules at the dendritic tip, which has been hypothesized to facilitate dendritic transport sustaining neuronal function [19].

To examine whether the neuronal centrosome contributes to either of the above processes, morphogenesis or dendritic trafficking, we took advantage of the tissue-specific degradation system described earlier. Depletion of SPD-5 early but not late in neuronal differentiation resulted in pronounced defects in the dyefill assay, which monitors uptake of the lipophilic dye DiI by surface exposed ciliated nerve endings [38] (Figure S3D). Defects were rescued by expression of a non-degradable form of SPD-5, confirming specificity of the phenotype. Conceptually, a dyefill phenotype could arise from a failure of dendrite extension or cilium assembly, or indeed a failure to form the neuron in the first place. An examination of the IFT marker CHE-11 in the phasmid neurons of SPD-5-depleted animals (Figure 3E, F) showed that terminal cell division was largely unaffected. However, we found evidence for the other two types of defects. Thus, neuronal morphology was clearly perturbed in a subset of animals, with collapsed dendrites indicating a failure of retrograde extension. Cilia were also frequently missing. While the majority of phasmids displayed both fully elongated dendrites and cilia with IFT occurring at roughly normal rates (Figure S3E, F), in many of those neurons intracellular trafficking was clearly perturbed, with CHE-11 particles accumulating along the dendrites (Figure 3E, F). The functional consequences for ciliogenesis could also be observed at the ultrastructural level, with transmission electron micrographs revealing missing axonemes as well as axonemes with less than 9 or incomplete doublet microtubules (Figure 3G, S3G). Thus, loss of the neuronal centrosome perturbs both morphogenesis and the trafficking that sustains cilium assembly and function.

### Distinct mechanisms of mitotic and non-mitotic PCM assembly

What is most remarkable about the neuronal centrosome is that it not only persists throughout the lifetime of the animal but new scaffold continues to assemble despite the absence of the key regulator PLK-1 and the degeneration of the centriole-derived basal body at its core. In contrast, the mitotic PCM requires centrioles for further scaffold incorporation as well as mechanical stability, while PLK-1 phosphorylation is essential for SPD-5 polymer formation as well as maintenance [2]. The *C. elegans* genome contains two further PLK1 homologs, PLK-2, which functions partially redundantly with PLK-1 in the germline and early embryo [2, 39, 40], and PLK-3, a putative pseudogene. We therefore considered the possibility that one of these paralogs sustains neuronal PCM incorporation. However, we were unable to detect PLK-2 at the ciliary base (not shown) and co-deletion of both *plk-2* and *plk-3* did not appreciably affect neuronal SPD-5 localization (Figure S4A). Treatment of worms with the specific inhibitor BI2536 which is expected to inhibit all PLK1 homologs [2, 41, 42] further did not affect the ability of SPD-5 to reassemble at the ciliary base following heat shock-induced ZIF-1-mediated degradation (Figure 4A, B). PLK-1 phosphorylation sites on SPD-5 have previously been mapped, with phosphorylation on S653 and S658 accounting for the stimulating effect of PLK-1 on SPD-5 oligomerization *in vitro* and *in vivo* [4]. Mutating these sites to alanine in the context of an RNAi-resistant endogenous promoter GFP transgene resulted in a failure of mitotic PCM expansion and significant embryonic lethality following depletion of the endogenous protein (Figure S4B, 0% viability, n=207 embryos *spd-5(RNAi)*; 99.7%, n=289 embryos *spd-5(RNAi)* WT rescue, 58.6%, n=186 embryos *spd-5(RNAi)* S653A S658A rescue). However, the same transgene was fully functional in reforming a PCM at the ciliary base following heat shock-induced degradation of both it and endogenous, wild-type SPD-5, the latter being prevented from neuronal expression by mutation of the X-box element in its promoter (Figure 4C, D). While we cannot exclude that another kinase drives SPD-5 oligomerization in ciliated neurons, the most straightforward explanation is that PCM assembly occurs via spontaneous self-assembly of SPD-5, as also observed *in vitro* [4]. Could this also be the case elsewhere? While most attention in the *C. elegans* early embryo has been on the dramatic expansion of the PCM in mitosis, a small amount of SPD-5 is already present at sperm-derived centrioles from soon after exit from meiosis II, contributing amongst other things to polarity establishment [43]. SPD-2, PLK-1 and AIR-1 are already present at this early stage. Yet, simultaneous inhibition of both Aurora A and PLK-1 with MLN8237 [44, 45] and BI2536, respectively, coupled with RNAi-mediated depletion of SPD-2 leave initial PCM recruitment largely unaffected (Figure 4E). Thus, the difference in PCM assembly mechanisms may be not so much between neuron and early embryo as between interphase/G0 and mitosis, with only the rapid expansion upon mitotic entry requiring PLK-1 activity (Figure S4C, D).

**Figure 4:**
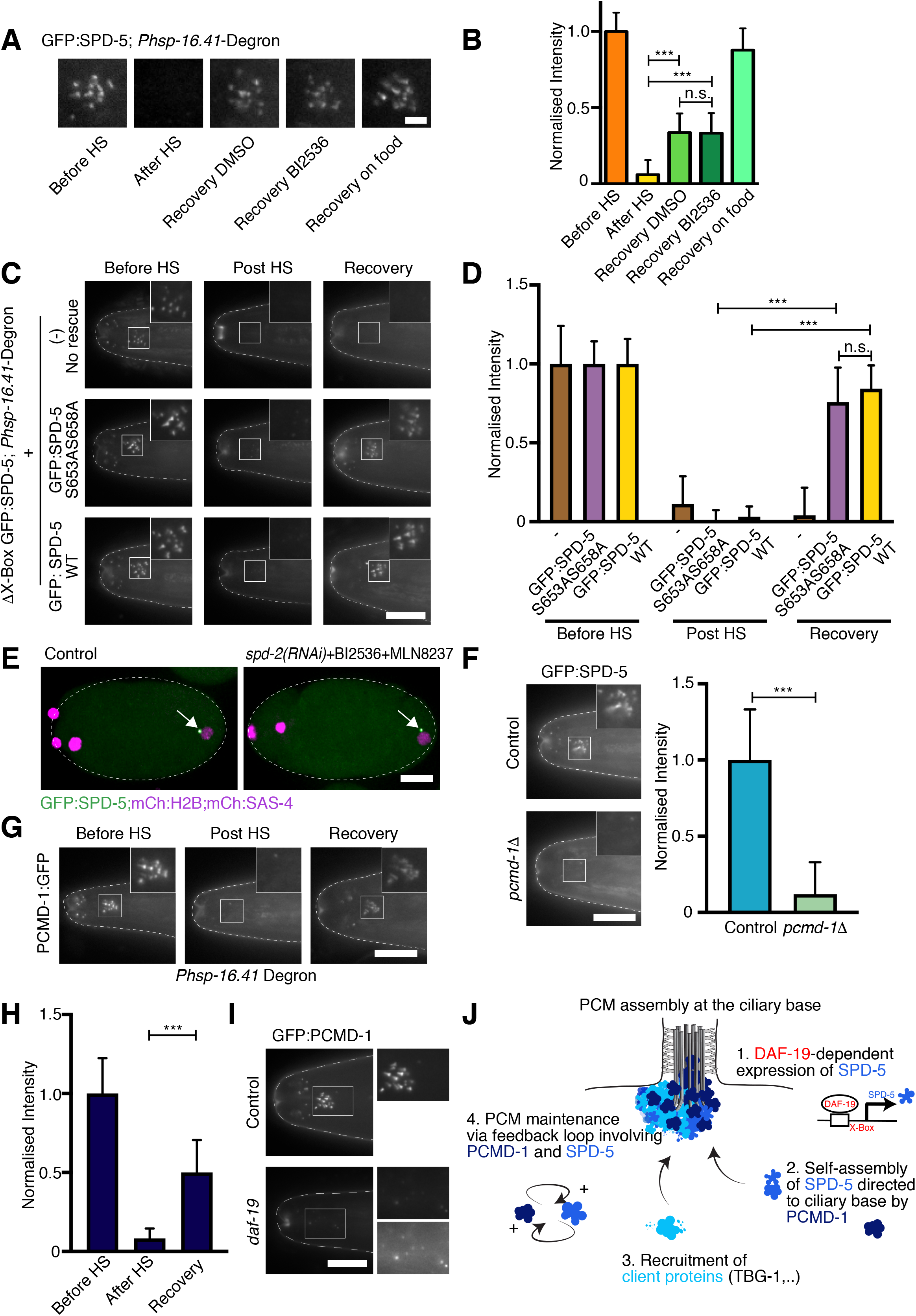
Distinct mechanisms of mitotic and non-mitotic PCM assembly. (**A**, **B**) PLK-1 inhibition does not affect *de novo* recruitment of SPD-5 following heat shock promoter-driven ZIF-mediated degradation. Images (A) and quantitation (B) of amphid GFP:SPD-5 signal before and immediately after heat shock, as well as after 24h of recovery in 40*μ*M BI2536 in M9, 2% DMSO in M9 (No food control) or on seeded NGM plates. Starvation reduces SPD-5 expression, but PLK1 inhibition does not further impair PCM scaffold assembly. n=22 before HS, 29 after HS, 24 recovery DMSO, 30 recovery BI2356, 27 recovery food. (**C**, **D**) SPD-5 PLK-1 phosphorylation mutant S653A S658A is capable of (self-) assembly at amphid ciliary base following heat shock promoter-driven degradation of endogenous (ΔX-box) GFP:SPD-5. Images (C) and quantitation (D) of GFP:SPD-5 signal for indicated strain genotypes before and after heat shock. Pre-heat shock signal independently normalized to 1 for each strain. n=10-16 animals no rescue, 15-17 rescue GFP:SPD-5 S653A S658A, 14-18 rescue GFP:SPD-5 WT. (**E**) Simultaneous inhibition of AIR-1 (MLN8237) and PLK-1 (BI2536) coupled with RNAi-mediated depletion of SPD-2 does not impair initial recruitment of PCM in the early embryo. Treated and untreated embryos co-expressing GFP:SPD-5, mCherry:SAS-4 and mCherry:H2B at equivalent stage of pronuclear expansion/ early S phase. (**F**) PCMD-1 is required for SPD-5 tethering to the ciliary base. GFP:SPD-5 signal in wild-type and *pcmd-1* mutant animals. n=17 animals control, 18 *pcmd-1* mutant. (**G**, **H**) Heat shock promoter-driven degradation shows PCMD-1 to be continuously expressed in amphid neurons. Images (G) and quantitation (H) of ciliary PCMD-1 signal before and after heat shock/recovery. n=13-16 animals per condition. (**I**) PCMD-1 signal is reduced in *daf-19* mutant animals but continues to form discrete foci. Lower inset panel auto-scaled for better presentation. Note dispersal of PCMD-1 foci due to widespread morphogenesis defects in *daf-19* mutants. (**J**) Model of PCM assembly at the ciliary base. SPD-5 is expressed under the daf-19 promoter and self-assembles at the ciliary base. Microtubule regulatory proteins concentrate within the SPD-5 PCM scaffold, creating an MTOC that contributes to cellular trafficking, neuronal morphogenesis and cilium assembly. PCMD-1 targets SPD-5 to the ciliary base. At the same time, the acentriolar PCM remains independent of ciliary structures, potentially through a feedback loop between PCMD-1 and SPD-5. Scale bars are 2*μ*m (A), 10*μ*m (others). Error bars are SD. *** t-test, P<0.001. ns. not significant. See also Figure S4.

### The pericentrin homolog PCMD-1 tethers SPD-5 to the ciliary base

The lack of centrioles as a core organizing structure presents another question: what maintains SPD-5 at the ciliary base and gives the PCM its structural integrity? The answer appears to lie in the recently described homolog of Pericentrin, PCMD-1, which plays a critical role in the initial recruitment of PCM to sperm-derived centrioles in the early embryo [7]. Like Pericentrin, PCMD-1 localizes close to the centriole barrel in the early embryo, with weaker signal extending into the surrounding PCM [7]. Unlike other centriolar proteins PCMD-1 is retained at the ciliary base. In both locations, fluorescence recovery after photobleaching shows the protein to be stably bound, with little exchange with the cytoplasmic pool (Figure S4E, F). Surprisingly, despite its important role in PCM recruitment in interphase, at least at low temperatures, subsequent mitotic PCM expansion and cell division is mostly unaffected by loss of PCMD-1, such that *pcmd-1* null mutants can be maintained in their homozygous state. At the L4 larval stage mutant animals display dyefill phenotypes similar to those following SPD-5 depletion (Figure S4G). An examination of SPD-5 localization reveals the reason why: a complete lack of PCM signal at the ciliary base (Figure 4F). ZIF-1 mediated degradation late in neuronal differentiation shows PCMD-1 loss also affects SPD-5 maintenance, albeit to a lesser extent (Figure S4H, I). PCMD-1 is therefore required to establish and maintain the interphase PCM in ciliated neurons. Consistent with an ongoing role of PCMD-1, PCMD-1 signal at the ciliary base recovers following heat shock-induced degradation, revealing ongoing neuronal expression (Figure 4G, H). How is this new protein targeted to the correct cellular location? We suggest this may be mediated by SPD-5, itself stably maintained at the ciliary base. There would then be a positive feedback loop between PCMD-1 and SPD-5, with PCMD-1 helping to recruit and stabilize the SPD-5 matrix, which in turn helps to recruit and maintain PCMD-1. While initially directed to occur around centrioles, such a feedback loop could stabilize the PCM following loss of its core organizing structure, the centriole. It would also explain why the PCM is robust to loss of specific ciliary structures, such as the IFT docking machinery in *hyls-1* mutants [6, 46] or the transition zone in *ccep-290;mksr-2;nphp-4* triple mutants [35] (Figure S4J), or indeed the entire ciliogenesis program in *daf-19* mutants, where residual SPD-5 and PCMD-1 inherited from prior cell cycles continue to form clear PCM foci despite the failure to form any ciliary structures [47] (Figure S2B, 4I).

In summary, we show that an acentriolar centrosome persists at the ciliary base in *C. elegans* sensory neurons. This centrosome is no mere remnant, but is actively maintained by continued expression of the scaffolding component SPD-5 under control of the RFX transcription factor DAF-19, contributing to neuronal morphogenesis and the cellular trafficking that sustains cilium assembly and maintenance (see model, Figure 4J). In contrast to mitotic PCM expansion, assembly and maintenance of the non-mitotic PCM, both in neurons and elsewhere in the worm, is independent of the mitotic regulator PLK-1. Instead, PCM assembly is likely driven by the self assembly properties of the scaffolding component SPD-5 [4]. At canonical centrosomes, centrioles act as the focal point for PCM assembly, while in differentiated tissues other cellular structures can assume that role [5]. Here, this is the ciliary base, which originally contained a centriole-derived basal body [6], but in other cases there is a transfer of PCM and consequently microtubule-organizing capacity to other cellular locations, such as the nuclear envelope in muscle [48] and the apical cell cortex in polarized epithelial cells [49]. A critical protein in maintaining the PCM scaffold at the ciliary base is Pericentrin/PCMD-1. PCMD-1 also tethers the growing PCM to centrioles in the early embryo [7], while Pericentrin helps establish a stable PCM scaffold at the nuclear envelope in muscle [48]. While much of the attention recently has been on the assembly of the PCM scaffold, formed by oligomerization of proteins such as Cnn/CDK5RAP2 and SPD-5 [4, 50], tethering proteins likely play a key role not only in targeting the PCM scaffold to specific cellular locations, but also in giving those microtubule organizing centers their requisite structural integrity [2].

## EXPERIMENTAL PROCEDURES

### *C. elegans* strains and culture conditions

*C. elegans* strains expressing endogenously tagged GFP:PCMD-1 [7], GFP:SPD-5, GFP:TAC-1 [2], GIP-2:GFP [26], PLK-1:GFP [51], SPD-2:GFP [13] TagRFP:GIP-1 [24], endogenous promoter-driven CHE-11:mKate, mCherry:AIR-1 [52], mCherry:SPD-2, GFP:SPD-5 [53], mCherry:HYLS-1 [35], MKS-6:mCherry [54], NOCA-1de:GFP, PTRN-1:GFP, TBG-1:GFP [26] and TBG-1:mCherry [4], *Ppie-1* promoter-driven mCherry:H2B [55], the *Pdyf-7* [30] and *Posm-6* [6] promoter-driven GFP nanobody-directed ZIF-1 degrons, and the loss of function alleles *plk-3(ok2812)* [56], *hyls-1(tm3067)* [22], *pcmd-1(syb975)* [7], *plk-2(ok1936)* [39], *ccep-290(tm4927)*, *mksr-2(tm2452)*, *nphp-4(tm925)* [35] and *daf-19(m86)* [31] have been described previously.

A strain with a mutated X-box sequence in the *spd-5* promoter was generated by Cas9-mediated homologous recombination using *in vitro*–synthesized Cas9-crRNA-tracrRNA complexes [57] with endogenously tagged GFP:SPD-5 as the starting point, replacing the original X-box motif (GTTTCC AT AGTAAC) with a reshuffled version unable to bind DAF-19 (TCTCGA GA TCATAT). The latter sequence also includes an XhoI restriction site (CTCGAG) facilitating identification of mutant animals. Endogenously tagged GFP:AIR-1 was similarly generated by Cas9-mediated homologous recombination using a plasmid-based protocol [58]. Successful integration was detected by single worm PCR (GFP:AIR-1)/ PCR and XhoI digest (ΔX-box GFP:SPD-5) and confirmed by PCR and sequencing. Strains expressing endogenous promoter-driven mNeonGreen:MZT-1, mCherry:SAS-4, GFP:SPD-5 (S653A S658A), mNeonGreenSPD-5 and GFP:ZYG-9 were generated by cloning the corresponding genomic locus including 5’ and 3’ regulatory sequences and N-terminal GFP/mNeonGreen/mCherry into the targeting vector pCFJ151 followed by Mos1-mediated transposition [59]/ gonad injection to form extrachromosomal arrays (GFP:ZYG-9, [60]). Both GFP:SPD-5 (S653A S658A) and the corresponding wild-type control [53] from which it is derived were rendered RNAi-resistant by re-encoding the sequence between nucleotides 500 to 1079 in the *spd-5* coding region to change the nucleotide sequence without altering the encoded amino acid sequence or codon usage [4]. For inducible degradation of GFP:PCMD-1 and GFP:SPD-5, strains expressing a GFP nanobody:ZIF-1 fusion [26], with and without mCherry:histone H2B as a visible marker coexpressed via an operon linker under the *hsp-16.41* heat shock promoter, were generated by Mos1 transposon insertion as above.

Dual color strains and strains carrying mutant alleles and ZIF-1 degrons were constructed by mating. The genotypes of all strains used are listed in the Key Resources Table. All strains were maintained at 23°C with the exception of strains carrying the temperature-sensitive allele *pcmd-1(syb975)*, which were kept at 16°C.

### Dye-fill assays

~100 worms were picked into 250*μ*l of M9 0.1% Triton, washed 3x with M9 and incubated for 1h in 0.2% DiI (Invitrogen) in M9. After incubation, worms were allowed to destain for at least 30 min on a seeded NGM plate before analysis. Dye accumulation in the cell bodies of amphid and phasmid neurons was scored on a Zeiss Axio Imager Z2 microscope equipped with a 63x 1.4NA Plan Apochromat objective, Lumencor SOLA SE II light source. For illustration purposes 1*μ*m z-series were acquired using a Photometrics CoolSNAP-HQ2 cooled CCD camera controlled by ZEN 2 blue software (Zeiss).

### Live imaging of embryos

Live imaging in early embryos was performed primarily on a Yokogawa CSU X1 spinning disk confocal mounted on a Zeiss Axio Observer Z1 inverted microscope equipped with a 63x 1.4NA Plan Apochromat objective, 120mW 405nm and 100mW 488nm and 561nm solid-state lasers, 2D-VisiFRAP Galvo FRAP module, Photometrics CoolSNAP-HQ2 cooled CCD and Hamamatsu ImagEM X2 EM-CCD cameras and controlled by VisiView software (Visitron Systems). Embryos were dissected in meiosis medium (60% Leibowitz L-15 media, 25μM Hepes pH7.4, 0.5% Inulin, 20% heat-inactivated fetal bovine serum) and filmed without compression [61]. Low laser illumination (max power of 0.32mW) was used to minimize photobleaching. For quantitative analysis of PCM recruitment, 12×1μm GFP/mCherry z-series as well as single plane DIC images were acquired at irregular intervals from nuclear envelope breakdown until cytokinesis onset. For the experiments presented in Figures 4E, S1C and 4B-D images were acquired using the EM-CCD camera with no binning. All other images were acquired using the CCD camera with 2×2 binning. Image stacks were imported into MetaMorph (Molecular Devices) for post-acquisition processing.

### Live imaging of amphid and phasmid neurons

Live imaging of amphid and phasmid neurons as well as of the embryos in Figure S1B was performed primarily using a 60x 1.42NA Plan Apochromat objective (for Figure 1C a 100x 1.4NA Uplan S Apochromat objective) on a DeltaVision 2 Ultra microscope, equipped with 7-Color SSI module and sCMOS camera and controlled by Acquire Ultra acquisition software (GE Healthcare). For the experiments presented in Figure S2A and S2B another DeltaVision microscope equipped with a CoolSNAP-HQ2 cooled CCD camera (Photometrics) but the same 60x objective and 7-Color SSI module was used. Worms were anesthetized using 10mM tetramisole in M9 for 5min before mounting on 2% agarose pads and imaging. 0.1-0.3*μ*m GFP/mCherry z-series as well as single plane DIC images were collected for the amphid and phasmid neurons closest to the objective. For analysis of CHE-11 IFT trafficking, single-plane mKate images of phasmid cilia in L4 larvae were acquired every half second. Image stacks were imported into Fiji for post-acquisition processing.

### Fluorescence recovery after photobleaching

To examine the dynamics of PCM proteins in early embryos, embryos in the first mitotic division expressing the GFP fusion of interest were followed from early prophase, with 6×1*μ*m z-series acquired at irregular intervals as described above. Photobleaching was performed in prometaphase using the galvanometer point scanner to target a region encompassing the centrosome with the 405nm laser at 10mW power and embryos imaged until completion of cytokinesis. Embryos were analyzed only if centrosome signal was completely eliminated throughout the entire z-volume.

To examine PCM dynamics in ciliated neurons, L4 stage worms were anesthetized using 10mM tetramisole and mounted on 2% agarose pads. Photobleaching was performed on phasmid neurons targeting one or both PCM foci at the ciliary base using the spinning disk confocal set-up described above. Before and after photobleaching images 6×0.5*μ*m z-series were acquired at irregular intervals using the 63x objective and CCD camera with 2×2 binning. As for early embryos, low laser illumination (max power of 0.32mW) was used to minimize photobleaching during acquisition. Pharyngeal pumping was monitored to ensure animals remained healthy during the ~10min following photobleaching.

Image stacks were imported into MetaMorph (Molecular Devices) for post-acquisition processing.

### RNA-mediated interference

RNAi experiments were performed by soaking [62] using the dsRNAs listed in the Key Resources Table. Standard RNAi conditions of 48h at 16°C were used for *spd-2* and *spd-5*. For egg shell permeabilization by *ptr-2(RNAi)*, recovery time was reduced to 24h at 20°C to minimize phenotypes beyond embryo permeability.

### Degron-mediated depletion

To deplete SPD-5 and PCMD-1 specifically in ciliated neurons, we employed the GFP nanobody directed ZIF-1-mediated protein degradation system [26] to target endogenously GFP-tagged SPD-5 and PCMD-1, expresssing the GFP nanobody::ZIF-1 fusion under the tissue-specific promoters *Pdyf-7* and *Posm-6*, active at the time of dendrite elongation (comma-stage) and after initiation of ciliogenesis (3-fold stage), respectively [6]. For inducible degradation at different developmental stages, ZIF-1 was expressed under the heat-shock-inducible promoter *Phsp-16.41*, activated by a temperature upshift to 30°C for 5h.

### Inhibitor experiments

To inhibit AIR-1 and PLK-1 in early embryos, *ptr-2(RNAi)* permeabilized embryos were dissected in meiosis medium (60% Leibowitz L-15 media, 25μM Hepes pH7.4, 0.5% Inulin, 20% heat-inactivated fetal bovine serum) containing 20μM MLN8237 (Aurora A inhibitor, Selleckchem) and 20μM BI2536 (PLK1 inhibitor, Axon MedChem). To inhibit PLK-1 in ciliated neurons following transient depletion of SPD-5, L2-stage larvae expressing *hsp-16.*41 promoter-driven GFP-nanobody::ZIF-1 and endogenously GFP-tagged SPD-5 were first heat-shocked for 5h at 30°C to degrade existing SPD-5 at the ciliary base. Worms were then either allowed to recover for 24h at 20°C (Control) or removed from the plate, washed with PBS and transferred with an eyelash-tool to a drop containing 40*μ*M BI2536 or 2% DMSO (No food control) in M9 and incubated in a humidified chamber for 24h at 20°C.

### Transmission electron microscopy

L4-stage worms were prepared by chemical fixation as previously described [63]. In brief, worms were fixed in 2.5% glutaraldehyde in cytoskeleton buffer (100 mM methyl ester sulfonate, 150 mM NaCl, 5 mM EGTA, 5 mM MgCl_2_, 5 mM glucose in ddH_2_O, pH 6.1) overnight at 4°C. Samples were then washed 3x in the same buffer and post-fixed for 30min in 0.5% osmium tetroxide in buffer, washed 3x in buffer and 1x in ddH_2_O. Finally, samples were dehydrated for 15min each in 40%, 60%, 80%, 2x in 95% and finally 3x in 100% acetone. Samples were embedded in Agar100 resin after fixation and dehydration. 70 nm serial sections were then prepared, post-stained with aqueous uranyl acetate and lead citrate and examined with a Morgagni 268D microscope (FEI) equipped with an 11-megapixel Morada CCD camera (Olympus-SIS) and operated at 80 kV.

### Image analysis

Quantification of PCM signal in early embryos was performed in Metamorph as previously described [13]. GFP signal was measured on single planes for both centrosomes at each time point. Two variable size concentric regions were drawn around each centrosome, a smaller one encompassing the centrosome, and a larger one including the surrounding cytoplasm as background. The integrated GFP intensity was then calculated by subtracting the mean fluorescence intensity in the area between the two boxes (mean background) from the mean intensity in the smaller box and multiplying by the area of the smaller box. To compensate for centrosome movement in z within the embryo, measurements for both centrosomes at each time point were averaged. All measurements are normalized to their respective controls.

Quantification of PCM signal at the ciliary base in amphids was performed in Fiji. GFP signal was measured on single planes with two concentric regions as described for early embryos, with the inner box encompassing the amphid ciliary cluster, the outer box the surrounding background, avoiding regions of high autofluorescence. The encompassing regions were drawn on the image plane with the highest signal intensity. All measurements are normalized to their respective controls. For the PLK-1 inhibition experiment, measurements were normalized to levels before heat-shock. GIP-1 Shannon Entropy was calculated in Python using the entropy function from the sci-kit image package.

For FRAP analysis in early embryos and phasmid ciliated neurons, GFP signal for the bleached centrosome was quantified on single planes and the integrated fluorescence intensity calculated as described above. Measurements taken after photobleaching were normalized to the mean intensity of measurements in the 30s preceding photobleaching. To account for additional PCM recruitment during centrosome maturation in early embryos, recruitment profiles were generated for unbleached embryos during the same cell cycle stage and the calculated signal increase subtracted from the measured FRAP recovery profiles. GraphPad Prism was then used to fit the data to the exponential equation Y=A*(1−exp(−k*X))+B where A is the mobile fraction, B is the background left after the bleach, and the half life t_1/2_=ln(0.5)/−k. R^2^ values are the correlation coefficient obtained by fitting experimental data to the model. Data points on the graphs are the mean of the normalized GFP intensity measurements collected during the 40s interval centered on that point.

For analysis of CHE-11 dynamics in phasmid cilia the time-lapse sequence of images was first corrected for sample drift using the Correct 3D Drift plugin in Fiji. Next, a segmented line with a width of 3 pixels was drawn along the cilium and the KymographBuilder plugin used to generate a kymograph. The velocity of CHE-11 IFT particles was then calculated from the slopes of individual lines on the kymograph. For IFT flux rate analysis, the number of particles passing through the proximal segment of the cilium was counted over the course of 30s.

### Statistical Analysis

All error bars are standard deviation unless otherwise indicated. To compare samples in a specific experiment, unpaired t-tests were conducted using GraphPad Prism. For entropy measurements in Figure S3C Welch’s correction was applied. *, **, *** represent P-Values of <0.05, 0.01 and 0.001, respectively. Tests are comparing indicated condition to control unless otherwise specified.

## KEY RESOURCES TABLE

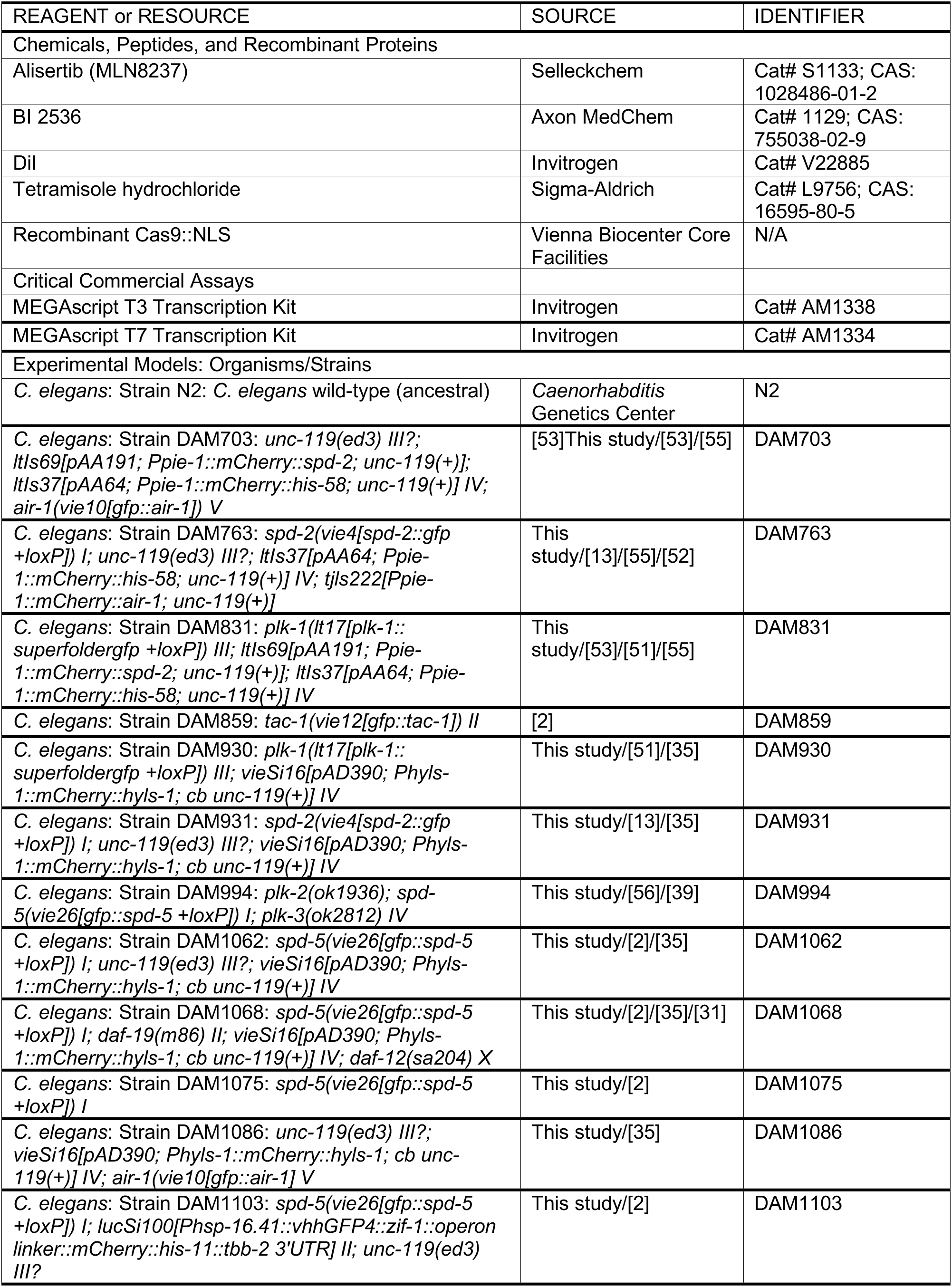
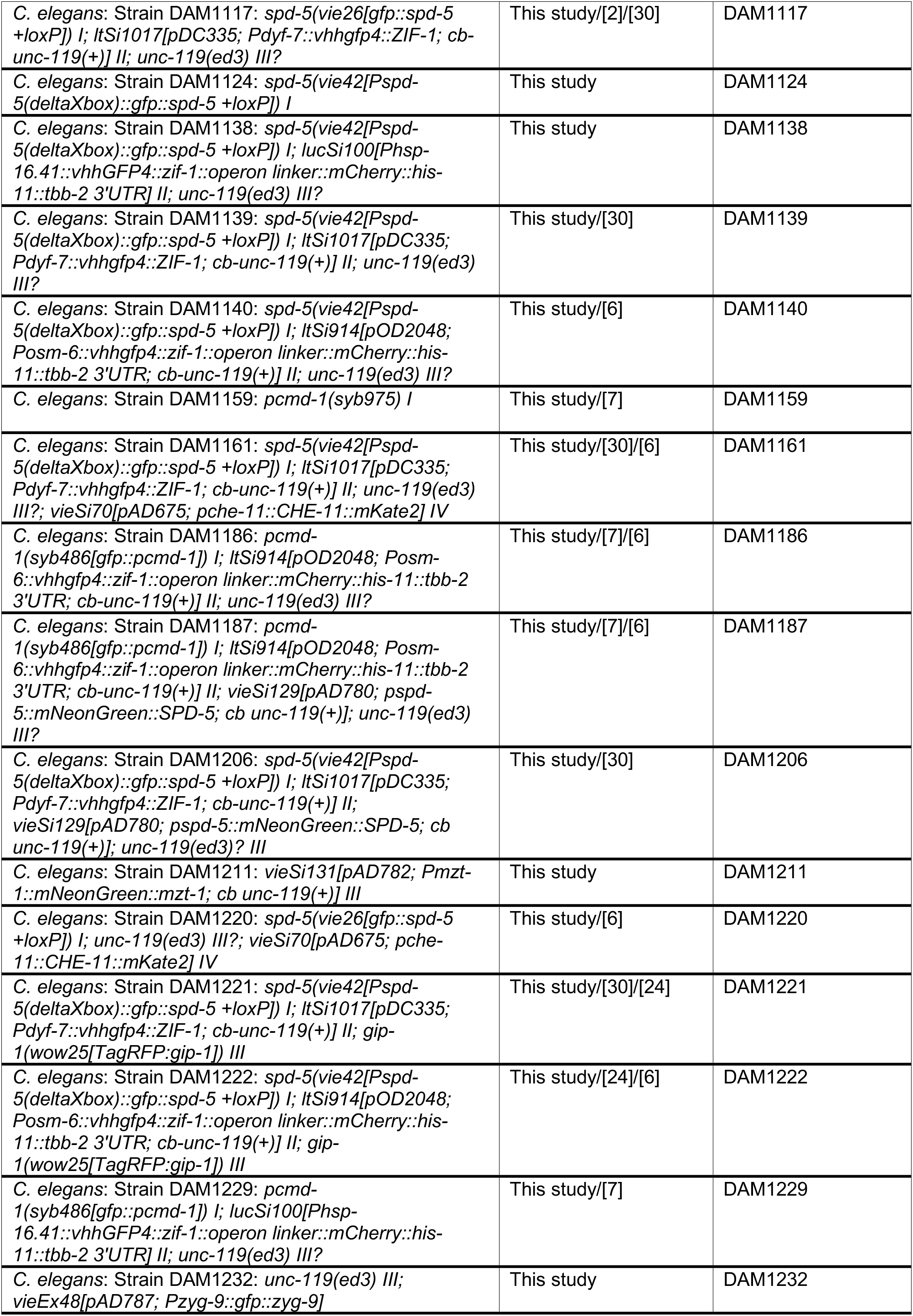
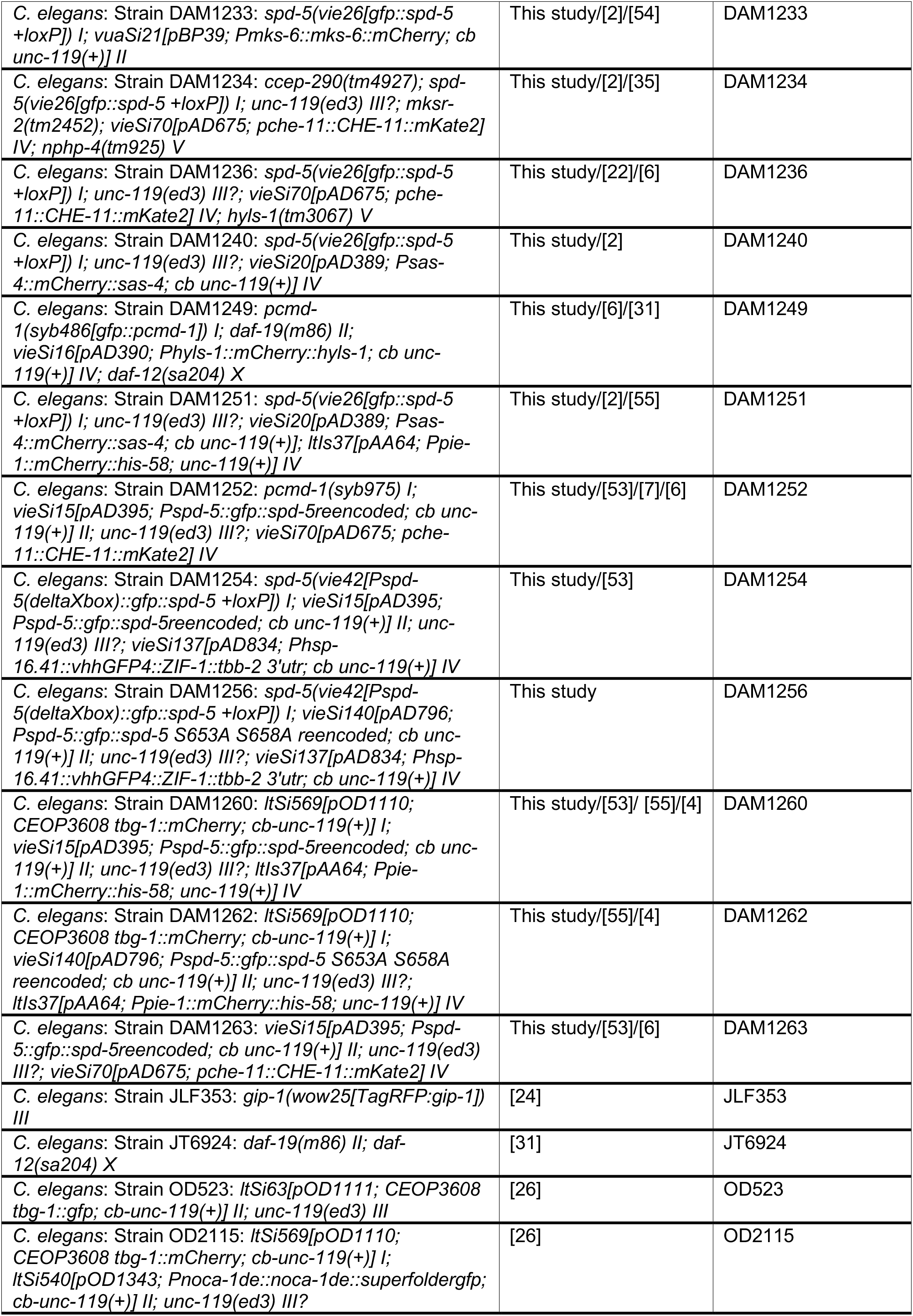
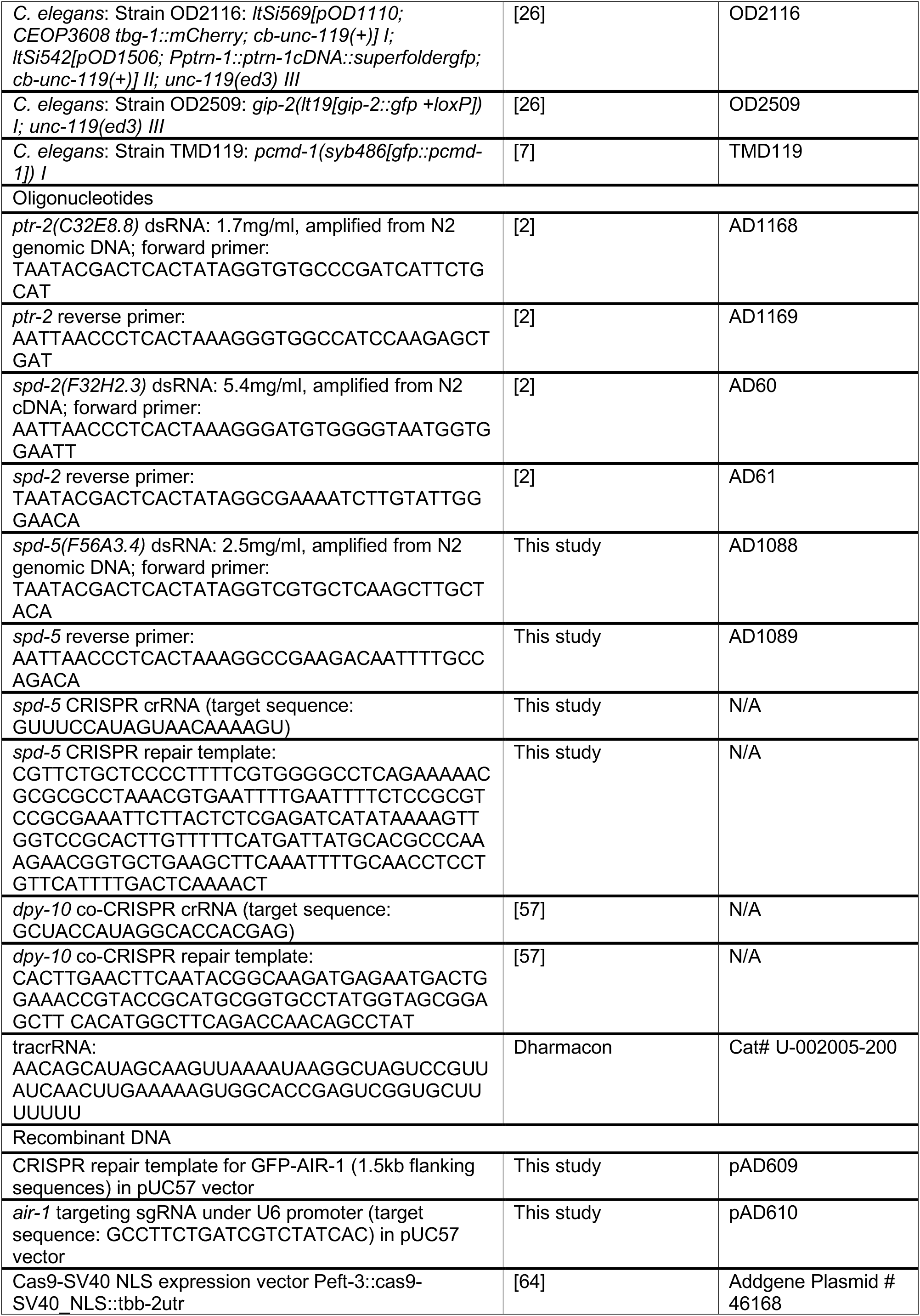
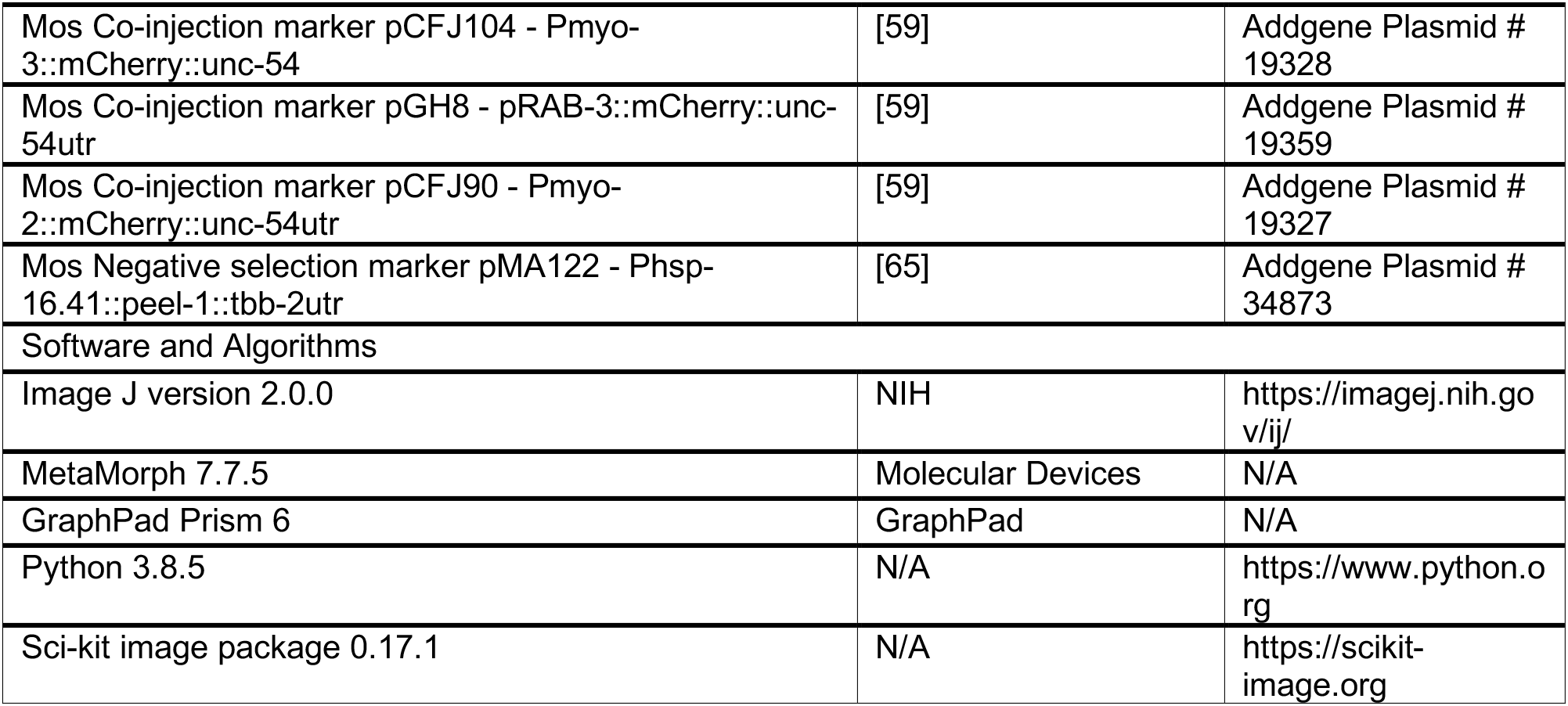

## ACKNOWLEDGEMENTS

We thank members of the Campbell and Dammermann lab for discussions, the *Caenorhabditis* Genetics Center, Dhanya Cheerambathur, Luisa Cochella, Jessica Feldman, Tamara Mikeladze-Dvali, Shohei Mitani, Karen Oegema, Erwin Peterman and Asako Sugimoto for strains and reagents, Christoph Sommer for help with data analysis, Nicole Drexler of the VBCF Electron Microscopy facility for help with EM, and Josef Gotzmann and Thomas Peterbauer of the Max Perutz Labs BioOptics facility for technical assistance. This work was supported by grant Y597-B20 from the Austrian Science Fund (FWF) to A.D..J. G. and T.L. are students of the DK Chromosome Dynamics, funded by the FWF (W1238-B20).

## AUTHOR CONTRIBUTIONS

J.G. and T.L. Conception and Design, Acquisition of Data, Analysis and Interpretation of Data, Drafting or Revising the Article; E.H. and M.D. Contributing Unpublished Essential Data or Reagents; A.D. Conception and Design, Analysis and Interpretation of Data, Drafting or Revising the Article.

## SUPPLEMENTAL FIGURE LEGENDS

**Figure S1:**
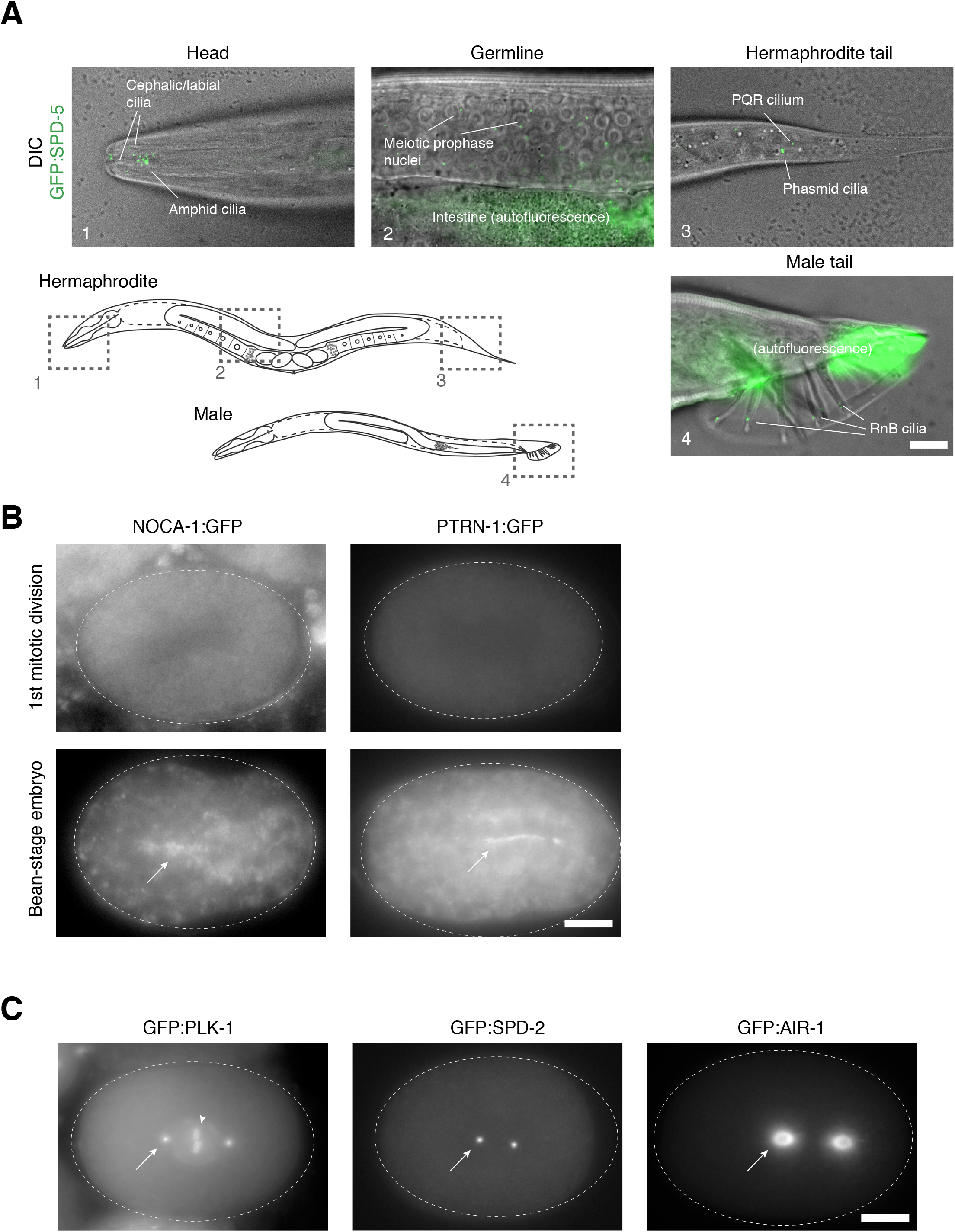
Further characterization of centrosomal protein localization. (**A**) Endogenously GFP-tagged SPD-5 localizes to the ciliary base in amphids, phasmids and other ciliated neurons in the head (cephalic, labial neurons) and tail (PQR, RnB neurons) of adult hermaphrodites and males. SPD-5 foci are also visible in the mitotic and meiotic germline, but there is otherwise little signal in the adult worm. Overlay of GFP foci on DIC images of the worm. Schematics adapted from WormBook <Zarkower, 2006 #1973>. (**B**) PTRN-1 and NOCA-1 prominently localize to the non-centrosomal microtubule-organizing center in the intestinal primordium of bean-stage embryos (~360min after fertilization) but not to mitotic centrosomes in the early embryo. (**C**) In contrast to the ciliary base (compare Figure 1E) there is prominent centrosomal signal for endogenously GFP-tagged PLK-1, SPD-2 and AIR-1 in mitotic embryos. SPD-2 and PLK-1 are highly concentrated at centrioles, while AIR-1 localizes to the PCM periphery and astral microtubules. PLK-1 also localizes to kinetochores (arrowhead). Scale bars are 10*μ*m.

**Figure S2:**
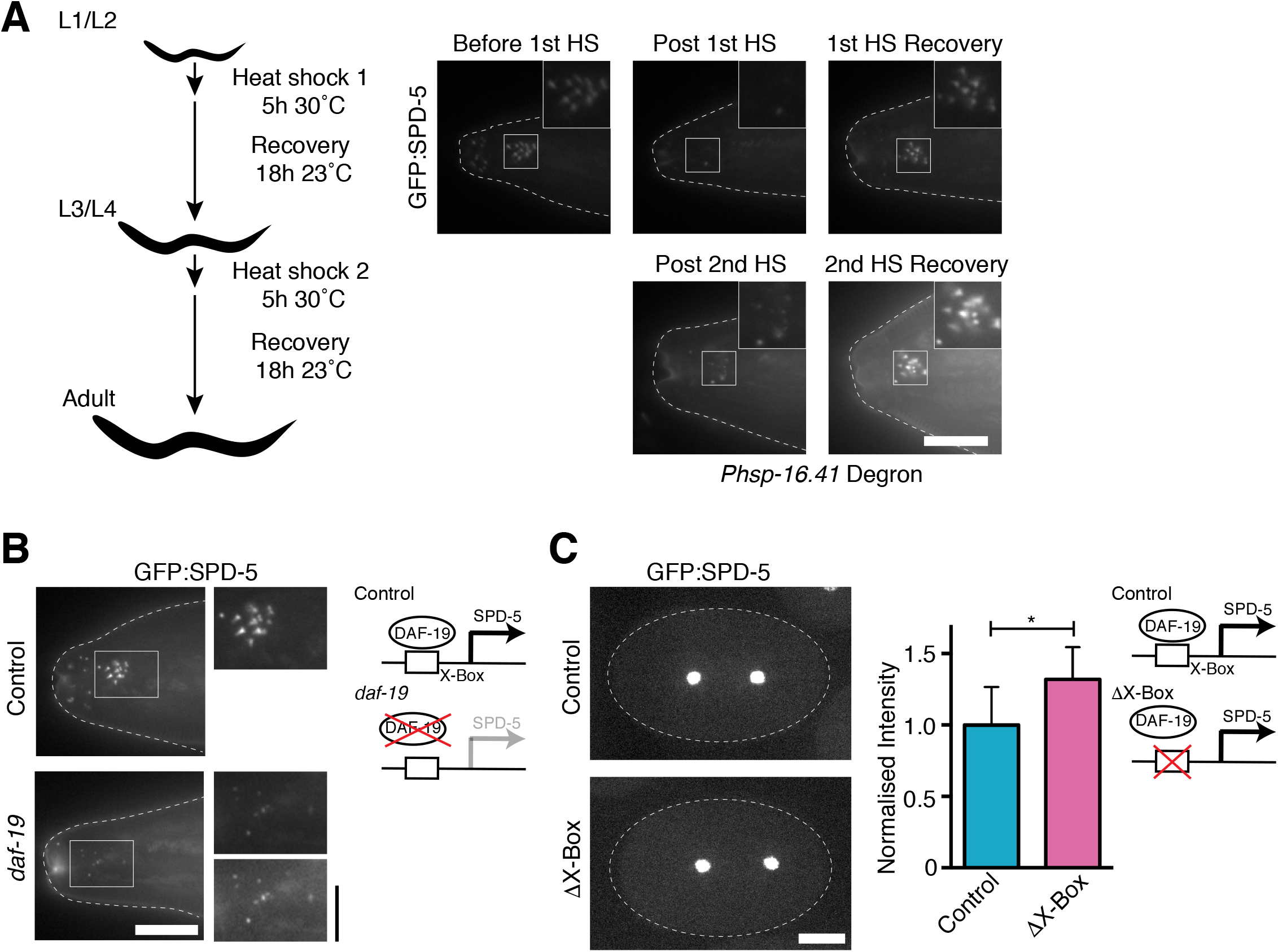
DAF-19 regulates SPD-5 expression in ciliated neurons. (**A**) GFP:SPD-5 signal continues to recover after repeated *Phsp-16.41* heat shock promoter-driven ZIF-mediated degradation, indicating ongoing expression. (**B**) GFP:SPD-5 signal is reduced in *daf-19* mutants. However, SPD-5 continues to form discrete foci despite failure of the ciliogenesis program. Lower inset panel auto-scaled for better presentation. Note dispersal of SPD-5 foci due to widespread morphogenesis defects in *daf-19* mutants. (**C**) Centrosomal GFP:SPD-5 signal in early embryos is unaffected (indeed marginally increased) in mutants lacking the DAF-19 X-box motif in the *spd-5* promoter. Signal normalized to metaphase controls. n=7 centrosomes control, 9 ΔX-box. Scale bars are 10*μ*m (A, B, C), 5 *μ*m (insets, B). Error bars are SD. * t-test, P<0.05.

**Figure S3:**
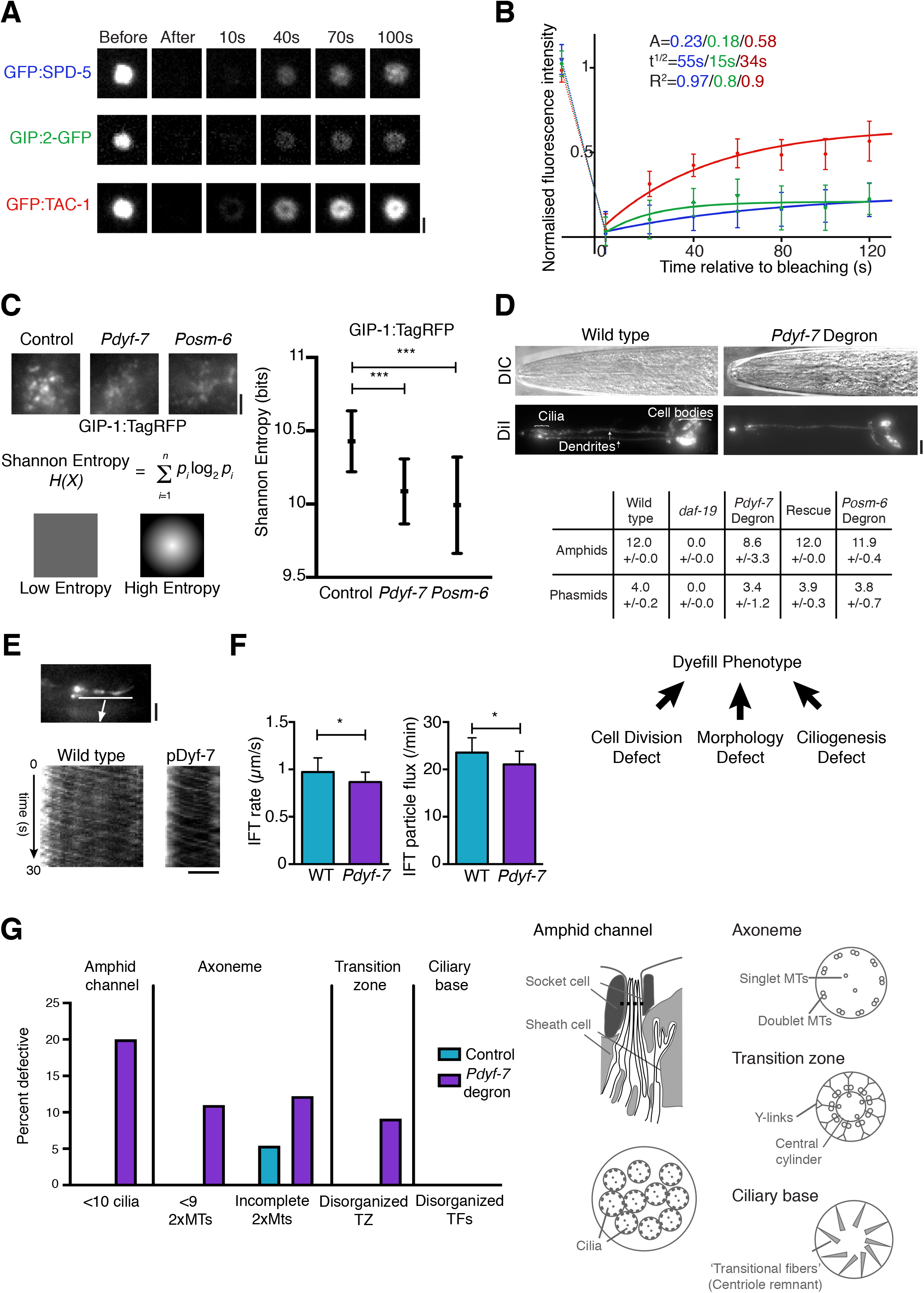
SPD-5 contributes to PCM scaffolding, neuronal morphogenesis and cilium assembly. (**A**, **B**) Fluorescence recovery after photobleaching (FRAP) analysis for PCM proteins in prometaphase-stage embryos. Images (A) and quantitations (B) for FRAP experiments performed on strains expressing GFP:SPD-5 (n=9 embryos), GIP-2:GFP (n=8) and GFP:TAC-1 (n=8). SPD-5 forms a PCM scaffold to which the γ-tubulin complex protein GIP-2 is stably bound, while TAC-1 remains in dynamic exchange with the cytoplasmic pool. Note that PCM continues to expand during centrosome maturation, resulting in apparent signal recovery for stable components. Quantitations are corrected for ongoing recruitment. (**C**) GIP-1:TagRFP PCM signal at the ciliary base is dispersed following degron-mediated depletion of SPD-5, as measured by Shannon entropy. Shannon Entropy was calculated based on the formula shown, where i represents states and p(i) the corresponding probability. Shannon entropy measures the average level of “information”, hence homogeneous intensity distributions have lower entropy values than non-homogeneous ones. n=19 amphids control, 23 *Pdyf-7* degron, 22 *Posm-6* degron. (**D**) Degron-mediated degradation of SPD-5 early (*Pdyf-7* promoter-driven) but not late (*Posm-6*) in neuronal differentiation results in a dye-fill phenotype, rescued by expression of non-degradable mNeonGreen:SPD-5. *daf-19* mutants shown as controls. The lipophilic dye DiI enters surface exposed ciliated neurons and accumulates in their cell bodies. A loss of dyefilling may reflect a failure of cell division, neuronal morphogenesis and/or cilium biogenesis. n>90 animals per condition. (**E**, **F**) Kymographs (E) and quantitations (F) for CHE-11:mKate2 particle movement within phasmid cilia following *Pdyf-7* promoter-driven degradation of SPD-5. IFT transport rates and particle flux appear largely unaffected by loss of neuronal PCM. n=9 cilia control, 19 *Pdyf-7* degron. (**G**) Quantitation of ultrastructural defects observed in transmission electron micrographs of control and *Pdyf-7* degron SPD-5-depleted animals shown in Figure 3G. Amphid channels in wild-type invariably contain 10 cilia. Likewise, the proximal segment of the axoneme and transition zones contain 9 doublet microtubules, a variable number of inner singlet microtubules and in the transition zone a central cylinder. 9 so-called ‘transitional fibers’ at the ciliary base represent the centriole remnant [6]. A number of these features are disrupted in SPD-5 depleted animals. 4 wild-type and 6 degron animals examined. n= 4/10 amphid channels, 37/82 axonemes, 16/22 transition zones and 9/18 ciliary bases. Phenotypes were scored only where sections provided sufficent ultrastructural detail. Scale bars are 2*μ*m (A, C, E), 10*μ*m (D). Error bars are 90% CI (B), SD (C, D, F). * t-test, P<0.05, ***P<0.001.

**Figure S4:**
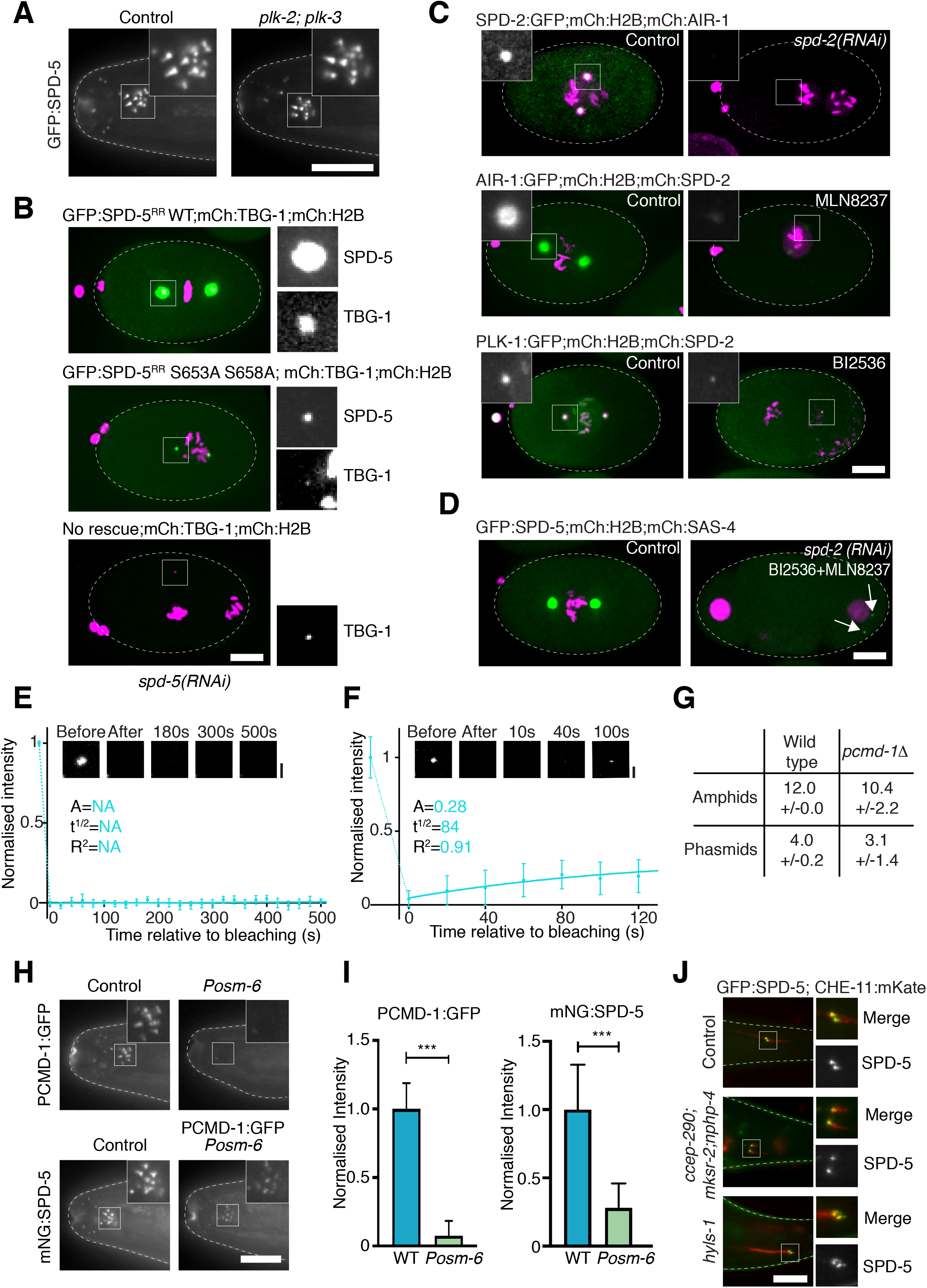
Further characterization of SPD-5 targeting and assembly at mitotic and non-mitotic centrosomes. (**A**) SPD-5 localization at the ciliary base is independent of *plk-2* and *plk-3*. Amphid GFP:SPD-5 signal in control and *plk-2*;*plk-3* mutant animals. (**B**) SPD-5 PLK-1 phosphorylation mutant S653A S658A is deficient in mitotic PCM assembly. Embryos expressing wild-type and mutant RNAi-resistant GFP:SPD-5 transgenes as well as mCherry:TBG-1 and mCherry:H2B depleted of endogenous SPD-5 by RNAi. (**C**) RNAi-mediated depletion of SPD-2 and treatment with inhibitors of AIR-1 (MLN8237) and PLK-1 (BI2536) impairs spindle assembly in early embryos. Images show prometaphase stage embryos in control and RNAi/inhibitor-treated conditions for each perturbation. Insets show targeted protein. *spd-2(RNAi)* eliminates all detectable SPD-2 signal. AIR-1 and PLK-1 inhibitors reduce but do not eliminate centrosomal targeting of the respective kinase. (**D**) Mitotic PCM expansion is abolished following simultaneous inhibition of AIR-1 (MLN8237) and PLK-1 (BI2536) coupled with RNAi-mediated depletion of SPD-2. Still images from time lapse sequences of treated and untreated embryos co-expressing GFP:SPD-5, mCherry:SAS-4 and mCherry:H2B at equivalent stage of development. (**E**, **F**) Fluorescence recovery after photobleaching (FRAP) analysis shows PCMD-1 to be stably incorporated at the ciliary base as well as at centrioles in early embryos. Images and quantitations for FRAP experiments performed in phasmid neurons (E, n=7 phasmids) and prometaphase-stage embryos (F, n=12). (**G**) *pcmd-1* mutants display dyefill defects comparable to those following SPD-5 depletion. Dyefill phenotypes for amphid and phasmid neurons in control and *pcmd-1Δ* mutants. n>90 animals per condition. (**H**) PCMD-1 is required for SPD-5 maintenance at the ciliary base. *Posm-6* degron-mediated depletion of PCMD-1 late in embryogenesis results in a loss of SPD-5 from the ciliary base. (**I**) Quantitation of PCMD-1 and SPD-5 signal from images as in (H). n=17 animals PCMD-1:GFP control, 18 Posm-6 degron, 24 animals mNeonGreen:SPD-5 control, 13 Posm-6 degron. (**J**) SPD-5 PCM signal is unaffected in mutants defective in transition zone assembly (*ccep-290;mksr-2;nphp-4* triple mutants) and in which the centriole remnant is destabilized *(hyls-1* mutants). Scale bars are 10*μ*m. Error bars are 90% CI (E, F), SD (I). *** t-test, P<0.001.

## REFERENCES

1. Conduit, P.T., Wainman, A., and Raff, J.W. (2015). Centrosome function and assembly in animal cells. Nat Rev Mol Cell Biol 16, 611–624.

2. Cabral, G., Laos, T., Dumont, J., and Dammermann, A. (2019). Differential Requirements for Centrioles in Mitotic Centrosome Growth and Maintenance. Dev Cell 50, 355–366.e356.

3. Feng, Z., Caballe, A., Wainman, A., Johnson, S., Haensele, A.F.M., Cottee, M.A., Conduit, P.T., Lea, S.M., and Raff, J.W. (2017). Structural Basis for Mitotic Centrosome Assembly in Flies. Cell 169, 1078–1089.e1013.

4. Woodruff, J.B., Wueseke, O., Viscardi, V., Mahamid, J., Ochoa, S.D., Bunkenborg, J., Widlund, P.O., Pozniakovsky, A., Zanin, E., Bahmanyar, S., et al. (2015). Centrosomes. Regulated assembly of a supramolecular centrosome scaffold in vitro. Science 348, 808–812.

5. Sanchez, A.D., and Feldman, J.L. (2017). Microtubule-organizing centers: from the centrosome to non-centrosomal sites. Curr Opin Cell Biol 44, 93–101.

6. Serwas, D., Su, T.Y., Roessler, M., Wang, S., and Dammermann, A. (2017). Centrioles initiate cilia assembly but are dispensable for maturation and maintenance in C. elegans. J Cell Biol 216, 1659–1671.

7. Erpf, A.C., Stenzel, L., Memar, N., Antoniolli, M., Osepashvili, M., Schnabel, R., Conradt, B., and Mikeladze-Dvali, T. (2019). PCMD-1 Organizes Centrosome Matrix Assembly in C. elegans. Curr Biol 29, 1324–1336.e1326.

8. Pintard, L., and Bowerman, B. (2019). Mitotic Cell Division in Caenorhabditis elegans. Genetics 211, 35–73.

9. Kitagawa, D., Vakonakis, I., Olieric, N., Hilbert, M., Keller, D., Olieric, V., Bortfeld, M., Erat, M.C., Fluckiger, I., Gonczy, P., et al. (2011). Structural basis of the 9-fold symmetry of centrioles. Cell 144, 364–375.

10. van Breugel, M., Hirono, M., Andreeva, A., Yanagisawa, H.A., Yamaguchi, S., Nakazawa, Y., Morgner, N., Petrovich, M., Ebong, I.O., Robinson, C.V., et al. (2011). Structures of SAS-6 suggest its organization in centrioles. Science 331, 1196–1199.

11. Woodruff, J.B., Ferreira Gomes, B., Widlund, P.O., Mahamid, J., Honigmann, A., and Hyman, A.A. (2017). The Centrosome Is a Selective Condensate that Nucleates Microtubules by Concentrating Tubulin. Cell 169, 1066–1077.e1010.

12. Hamill, D.R., Severson, A.F., Carter, J.C., and Bowerman, B. (2002). Centrosome maturation and mitotic spindle assembly in C. elegans require SPD-5, a protein with multiple coiled-coil domains. Dev Cell 3, 673–684.

13. Laos, T., Cabral, G., and Dammermann, A. (2015). Isotropic incorporation of SPD-5 underlies centrosome assembly in C. elegans. Curr Biol 25, R648–R649.

14. Enos, S.J., Dressler, M., Gomes, B.F., Hyman, A.A., and Woodruff, J.B. (2018). Phosphatase PP2A and microtubule-mediated pulling forces disassemble centrosomes during mitotic exit. Biol Open 7.

15. Sulston, J.E., Schierenberg, E., White, J.G., and Thomson, J.N. (1983). The embryonic cell lineage of the nematode Caenorhabditis elegans. Dev Biol 100, 64–119.

16. Lu, Y., and Roy, R. (2014). Centrosome/Cell Cycle Uncoupling and Elimination in the Endoreduplicating Intestinal Cells of *C. elegans*. PLoS One 9, e110958.

17. Nechipurenko, I.V., Berciu, C., Sengupta, P., and Nicastro, D. (2017). Centriolar remodeling underlies basal body maturation during ciliogenesis in Caenorhabditis elegans. eLife 6.

18. Bobinnec, Y., Fukuda, M., and Nishida, E. (2000). Identification and characterization of Caenorhabditis elegans gamma-tubulin in dividing cells and differentiated tissues. J Cell Sci 113 Pt 21, 3747–3759.

19. Harterink, M., Edwards, S.L., de Haan, B., Yau, K.W., van den Heuvel, S., Kapitein, L.C., Miller, K.G., and Hoogenraad, C.C. (2018). Local microtubule organization promotes cargo transport in C. elegans dendrites. J Cell Sci 131.

20. Wiese, C., and Zheng, Y. (2000). A new function for the gamma-tubulin ring complex as a microtubule minus-end cap. Nat Cell Biol 2, 358–364.

21. Inglis, P.N., Ou, G., Leroux, M.R., and Scholey, J.M. (2007). The sensory cilia of Caenorhabditis elegans. WormBook, 1–22.

22. Dammermann, A., Pemble, H., Mitchell, B.J., McLeod, I., Yates, J.R., 3rd, Kintner, C., Desai, A.B., and Oegema, K. (2009). The hydrolethalus syndrome protein HYLS-1 links core centriole structure to cilia formation. Genes Dev 23, 2046–2059.

23. Hannak, E., Oegema, K., Kirkham, M., Gonczy, P., Habermann, B., and Hyman, A.A. (2002). The kinetically dominant assembly pathway for centrosomal asters in Caenorhabditis elegans is gamma-tubulin dependent. J Cell Biol 157, 591–602.

24. Sallee, M.D., Zonka, J.C., Skokan, T.D., Raftrey, B.C., and Feldman, J.L. (2018). Tissue-specific degradation of essential centrosome components reveals distinct microtubule populations at microtubule organizing centers. PLoS biology 16, e2005189.

25. Srayko, M., Quintin, S., Schwager, A., and Hyman, A.A. (2003). Caenorhabditis elegans TAC-1 and ZYG-9 form a complex that is essential for long astral and spindle microtubules. Curr Biol 13, 1506–1511.

26. Wang, S., Wu, D., Quintin, S., Green, R.A., Cheerambathur, D.K., Ochoa, S.D., Desai, A., and Oegema, K. (2015). NOCA-1 functions with gamma-tubulin and in parallel to Patronin to assemble non-centrosomal microtubule arrays in C. elegans. eLife 4, e08649.

27. Kemp, C.A., Kopish, K.R., Zipperlen, P., Ahringer, J., and O’Connell, K.F. (2004). Centrosome Maturation and Duplication in C. elegans Require the Coiled-Coil Protein SPD-2. Dev Cell 6, 511–523.

28. Pelletier, L., Özlü, N., Hannak, E., Cowan, C., Habermann, B., Ruer, M., Müller-Reichert, T., and Hyman, A.A. (2004). The Caenorhabditis elegans Centrosomal Protein SPD-2 Is Required for both Pericentriolar Material Recruitment and Centriole Duplication. Curr Biol 14, 863–873.

29. Hannak, E., Kirkham, M., Hyman, A.A., and Oegema, K. (2001). Aurora-A kinase is required for centrosome maturation in Caenorhabditis elegans. J Cell Biol 155, 1109–1116.

30. Cheerambathur, D.K., Prevo, B., Chow, T.L., Hattersley, N., Wang, S., Zhao, Z., Kim, T., Gerson-Gurwitz, A., Oegema, K., Green, R., et al. (2019). The Kinetochore-Microtubule Coupling Machinery Is Repurposed in Sensory Nervous System Morphogenesis. Dev Cell 48, 864–872.e867.

31. Swoboda, P., Adler, H.T., and Thomas, J.H. (2000). The RFX-type transcription factor DAF-19 regulates sensory neuron cilium formation in C. elegans. Mol Cell 5, 411–421.

32. Efimenko, E., Bubb, K., Mak, H.Y., Holzman, T., Leroux, M.R., Ruvkun, G., Thomas, J.H., and Swoboda, P. (2005). Analysis of xbx genes in C. elegans. Development 132, 1923–1934.

33. Tavernarakis, N., Wang, S.L., Dorovkov, M., Ryazanov, A., and Driscoll, M. (2000). Heritable and inducible genetic interference by double-stranded RNA encoded by transgenes. Nat Genet 24, 180–183.

34. Heiman, M.G., and Shaham, S. (2009). DEX-1 and DYF-7 establish sensory dendrite length by anchoring dendritic tips during cell migration. Cell 137, 344–355.

35. Schouteden, C., Serwas, D., Palfy, M., and Dammermann, A. (2015). The ciliary transition zone functions in cell adhesion but is dispensable for axoneme assembly in C. elegans. J Cell Biol 210, 35–44.

36. Kirszenblat, L., Pattabiraman, D., and Hilliard, M.A. (2011). LIN-44/Wnt directs dendrite outgrowth through LIN-17/Frizzled in C. elegans Neurons. PLoS biology 9, e1001157.

37. Baas, P.W., and Lin, S. (2011). Hooks and comets: The story of microtubule polarity orientation in the neuron. Developmental neurobiology 71, 403–418.

38. Hedgecock, E.M., Culotti, J.G., Thomson, J.N., and Perkins, L.A. (1985). Axonal guidance mutants of Caenorhabditis elegans identified by filling sensory neurons with fluorescein dyes. Dev Biol 111, 158–170.

39. Labella, S., Woglar, A., Jantsch, V., and Zetka, M. (2011). Polo Kinases Establish Links between Meiotic Chromosomes and Cytoskeletal Forces Essential for Homolog Pairing. Developmental Cell 21, 948–958.

40. Nishi, Y., Rogers, E., Robertson, S.M., and Lin, R. (2008). Polo kinases regulate C. elegans embryonic polarity via binding to DYRK2-primed MEX-5 and MEX-6. Development 135, 687–697.

41. Lenart, P., Petronczki, M., Steegmaier, M., Di Fiore, B., Lipp, J.J., Hoffmann, M., Rettig, W.J., Kraut, N., and Peters, J.M. (2007). The small-molecule inhibitor BI 2536 reveals novel insights into mitotic roles of polo-like kinase 1. Curr Biol 17, 304–315.

42. Noatynska, A., Panbianco, C., and Gotta, M. (2010). SPAT-1/Bora acts with Polo-like kinase 1 to regulate PAR polarity and cell cycle progression. Development 137, 3315–3325.

43. Cowan, C.R., and Hyman, A.A. (2004). Centrosomes direct cell polarity independently of microtubule assembly in C. elegans embryos. Nature 431, 92–96.

44. Manfredi, M.G., Ecsedy, J.A., Chakravarty, A., Silverman, L., Zhang, M., Hoar, K.M., Stroud, S.G., Chen, W., Shinde, V., Huck, J.J., et al. (2011). Characterization of Alisertib (MLN8237), an investigational small-molecule inhibitor of aurora A kinase using novel in vivo pharmacodynamic assays. Clinical cancer research : an official journal of the American Association for Cancer Research 17, 7614–7624.

45. Sumiyoshi, E., Fukata, Y., Namai, S., and Sugimoto, A. (2015). Caenorhabditis elegans Aurora A kinase is required for the formation of spindle microtubules in female meiosis. Mol Biol Cell 26, 4187–4196.

46. Wei, Q., Zhang, Y., Schouteden, C., Zhang, Y., Zhang, Q., Dong, J., Wonesch, V., Ling, K., Dammermann, A., and Hu, J. (2016). The hydrolethalus syndrome protein HYLS-1 regulates formation of the ciliary gate. Nature communications 7, 12437.

47. Perkins, L.A., Hedgecock, E.M., Thomson, J.N., and Culotti, J.G. (1986). Mutant sensory cilia in the nematode Caenorhabditis elegans. Dev Biol 117, 456–487.

48. Srsen, V., Fant, X., Heald, R., Rabouille, C., and Merdes, A. (2009). Centrosome proteins form an insoluble perinuclear matrix during muscle cell differentiation. BMC Cell Biol 10, 28.

49. Feldman, J.L., and Priess, J.R. (2012). A role for the centrosome and PAR-3 in the hand-off of MTOC function during epithelial polarization. Curr Biol 22, 575–582.

50. Conduit, P.T., Feng, Z., Richens, J.H., Baumbach, J., Wainman, A., Bakshi, S.D., Dobbelaere, J., Johnson, S., Lea, S.M., and Raff, J.W. (2014). The centrosome-specific phosphorylation of Cnn by Polo/Plk1 drives Cnn scaffold assembly and centrosome maturation. Dev Cell 28, 659–669.

51. Martino, L., Morchoisne-Bolhy, S., Cheerambathur, D.K., Van Hove, L., Dumont, J., Joly, N., Desai, A., Doye, V., and Pintard, L. (2017). Channel Nucleoporins Recruit PLK-1 to Nuclear Pore Complexes to Direct Nuclear Envelope Breakdown in C. elegans. Dev Cell 43, 157–171.e157.

52. Toya, M., Iida, Y., and Sugimoto, A. (2010). Imaging of mitotic spindle dynamics in Caenorhabditis elegans embryos. Methods Cell Biol 97, 359–372.

53. Cabral, G., Sanegre Sans, S., Cowan, C.R., and Dammermann, A. (2013). Multiple Mechanisms Contribute to Centriole Separation in C. elegans. Curr Biol 23, 1380–1387.

54. Prevo, B., Mangeol, P., Oswald, F., Scholey, J.M., and Peterman, E.J. (2015). Functional differentiation of cooperating kinesin-2 motors orchestrates cargo import and transport in C. elegans cilia. Nat Cell Biol 17, 1536–1545.

55. McNally, K., Audhya, A., Oegema, K., and McNally, F.J. (2006). Katanin controls mitotic and meiotic spindle length. J Cell Biol 175, 881–891.

56. Barber, N.C. (2011). Regulation of meiotic chromosome interactions in Caenorhabditis elegans, (University of California, Berkeley).

57. Paix, A., Folkmann, A., Rasoloson, D., and Seydoux, G. (2015). High Efficiency, Homology-Directed Genome Editing in Caenorhabditis elegans Using CRISPR-Cas9 Ribonucleoprotein Complexes. Genetics 201, 47–54.

58. Dickinson, D.J., Ward, J.D., Reiner, D.J., and Goldstein, B. (2013). Engineering the Caenorhabditis elegans genome using Cas9-triggered homologous recombination. Nat Methods 10, 1028–1034.

59. Frokjaer-Jensen, C., Davis, M.W., Hopkins, C.E., Newman, B.J., Thummel, J.M., Olesen, S.P., Grunnet, M., and Jorgensen, E.M. (2008). Single-copy insertion of transgenes in Caenorhabditis elegans. Nat Genet 40, 1375–1383.

60. Mello, C.C., Kramer, J.M., Stinchcomb, D., and Ambros, V. (1991). Efficient gene transfer in C.elegans: extrachromosomal maintenance and integration of transforming sequences. Embo j 10, 3959–3970.

61. Monen, J., Maddox, P.S., Hyndman, F., Oegema, K., and Desai, A. (2005). Differential role of CENP-A in the segregation of holocentric C. elegans chromosomes during meiosis and mitosis. Nat Cell Biol 7, 1248–1255.

62. Green, R.A., Kao, H.L., Audhya, A., Arur, S., Mayers, J.R., Fridolfsson, H.N., Schulman, M., Schloissnig, S., Niessen, S., Laband, K., et al. (2011). A high-resolution C. elegans essential gene network based on phenotypic profiling of a complex tissue. Cell 145, 470–482.

63. Serwas, D., and Dammermann, A. (2015). Ultrastructural analysis of Caenorhabditis elegans cilia. Methods Cell Biol 129, 341–367.

64. Friedland, A.E., Tzur, Y.B., Esvelt, K.M., Colaiacovo, M.P., Church, G.M., and Calarco, J.A. (2013). Heritable genome editing in C. elegans via a CRISPR-Cas9 system. Nat Methods 10, 741–743.

65. Frokjaer-Jensen, C., Davis, M.W., Ailion, M., and Jorgensen, E.M. (2012). Improved Mos1-mediated transgenesis in C. elegans. Nat Methods 9, 117–118.

